# Frequency-specific changes in prefrontal activity associated with maladaptive belief updating in volatile environments in euthymic bipolar disorder

**DOI:** 10.1101/2024.03.26.586656

**Authors:** Marina Ivanova, Ksenia Germanova, Dmitry S. Petelin, Aynur Ragymova, Grigory Kopytin, Beatrice A. Volel, Vadim V. Nikulin, Maria Herrojo Ruiz

## Abstract

Bipolar disorder (BD) involves altered reward processing and decision-making, with inconsistencies across studies. Here, we integrated hierarchical Bayesian modelling with magnetoencephalography (MEG) to characterise maladaptive belief updating in this condition. First, we determined if previously reported increased learning rates in BD stem from a heightened expectation of environmental changes. Additionally, we examined if this increased expectation speeds up belief updating in decision-making, associated with modulation of rhythmic neural activity within the prefrontal, orbitofrontal, and anterior cingulate cortex (PFC, OFC, ACC). Twenty-two 22 euthymic BD and 27 healthy control (HC) participants completed a reward-based motor decision-making task in a volatile setting. Hierarchical Bayesian modelling revealed BD participants anticipated greater environmental volatility, resulting in a more stochastic mapping from beliefs to actions and paralleled by lower win rates and a reduced tendency to repeat rewarded actions than HC. Despite this, BD individuals adjusted their expectations of action-outcome contingencies more slowly, but both groups invigorated their actions similarly. On a neural level, while healthy individuals exhibited an alpha-beta suppression and gamma increase during belief updating, BD participants showed dampened effects, extending across the PFC, OFC, and ACC regions. This was accompanied by an abnormally increased beta-band directed information flow in BD. Overall, the results suggest euthymic BD individuals anticipate environmental change without adequately learning from it, contributing to maladaptive belief updating. Alterations in frequency-domain amplitude and functional connectivity within the PFC, OFC, and ACC during belief updating underlie the computational effects and could serve as potential indicators for predicting relapse in future research.

## Introduction

Bipolar disorder (BD) is a chronic affective condition characterised by episodes of elation, depression, and mixed states, interspersed with periods of clinical remission[1,2]. Alterations in reward processing and impaired decision-making performance have been associated with the condition[3,4], pointing to disrupted functional connectivity between the prefrontal cortex (PFC) and the mesolimbic reward system[5,6]. Yet findings across studies are variable, even when considering euthymic periods alone[5–7]. BD research has reported both heightened sensitivity to negative feedback and decreased learning from rewards, or the reverse[8,10–13]. Recently, Ossola et al.[14] found that in euthymic BD, attenuated belief updating from positive feedback forecasts relapse, highlighting the importance of investigating dynamic belief updating during euthymia.

Influential proposals advocate for the application of computational models in BD research to understand fluctuations in mood and reward processing[15–17]. These models, building upon previous neurocomputational work on mood instability[18], suggest that altered affective reactivity to reward and punishment in BD may elevate learning rates, even during euthymic phases, predisposing individuals to form stronger expectations about rewards or punishments. New empirical work supports this, revealing a tendency in BD for reward perception to be biased by fluctuations in the momentum of recent reward prediction errors[19]. An increased learning rate could also reflect a heightened anticipation of environmental changes in BD[15]. Indeed, seminal modelling frameworks support that agents learn faster when anticipating more frequent transitions in the environment[20,21]. However, the extent to which individuals with BD perceive their environment as more volatile and how this perception influences their belief updating and decision-making remains unexplored.

This study investigates the computational processes underlying alterations in decision-making in euthymic bipolar patients, compared to healthy participants, as they undertake a probabilistic reward-based learning task in a volatile environment. Our primary hypothesis is that individuals with euthymic BD overestimate environmental volatility, thereby increasing their learning about the statistical correlations within their environment. Such inflated volatility estimates could introduce ’noise’ into the decision-making process[22], leading to incorrect decisions. Alternatively, the reported deficits in decision-making performance during euthymic BD[3,4] could be explained by slower belief updating, aligning with recent empirical observations in valence-dependent learning[13].

To test these hypotheses, we employ the Hierarchical Gaussian Filter (HGF), a validated modelling framework based on Hierarchical Bayesian inference that describes individual learning dynamics in environments characterised by uncertainty and volatility[23–26]. We used the HGF to model how input about probabilistic reward outcomes and their change over time is integrated with prior beliefs during learning, resulting in posterior beliefs about the hidden states causing the observed outcomes[23,24]. Belief updates in the HGF are driven by prediction errors (PE)—the discrepancy between predictions and outcomes—and are modulated by the precision weights, where precision is defined as the inverse variance of belief distributions. This computational framework has already proven useful for understanding psychiatric conditions[13,27,28], and could offer insights into how affective states and mood dynamically shape adaptive learning[29–31].

To gain a more mechanistic understanding of the processes underlying the hypothesised computational alterations in euthymic BD, we additionally investigated the neural correlates of hierarchical belief updating using magnetoencephalography (MEG). Existing research supports the role of cortical oscillations in maintaining predictions and encoding PEs[32–35]. Specific frequency rhythms such as alpha (8-12 Hz) and beta (13-30 Hz) oscillations have been associated with the transmission of top-down predictions, and encoding precision, while gamma-band activity (> 30 Hz) has been linked to the propagation of PEs and precision-weighted PEs, pwPEs[33,35–37]. Importantly, disruptions in these rhythms are suggested to contribute to learning deficits observed in various psychiatric conditions, including anxiety, schizophrenia, and autism[37–39].

On a neural level, we hypothesised that biases in probabilistic reward-based learning in BD in a volatile setting can be reflected in alpha, beta, and gamma activity during the encoding of pwPE and precision. These alterations are expected to manifest in the orchestrated activity across decision-making brain areas, such as the prefrontal, anterior cingulate, and orbitofrontal cortex (PFC, ACC, OFC). These regions are involved in learning in volatile and uncertain settings[20,37,40,41], and form part of the fronto-striatal reward circuit, which exhibits disturbed connectivity in BD[9,10]. We therefore additionally hypothesised that changes in frequency-domain connectivity patterns between these regions during belief updating would occur in BD relative to healthy control participants.

Lastly, we aimed to determine whether the computations underlying decision-making deficits in euthymic BD influence the motivational aspects associated with the invigoration of movements. Evidence suggests that reward expectations can speed motor performance[42,43]. The nigrostriatal dopamine pathway, central to the ’dopamine hypothesis’ of BD[44,45], is crucial for invigorating future movements[46]. Moreover, individuals with BD have been shown to experience increases in energy and effort following success, thus demonstrating enhanced motor vigour effects[4,47]. Consequently, our final complementary hypothesis posits that the strength of predictions about reward contingencies in the environment will speed decision-related movements in euthymic BD more than in healthy individuals[48].

## Methods and Materials

### Participants

Participants included 22 bipolar patients (mean age: 29.1 years [SEM = 1.67], 17 females; **Table 1**), and 27 healthy participants (27.5 years [SEM = 1.18], 15 females). Bipolar participants were assessed by a consultant psychiatrist who confirmed the diagnosis of BD (I or II) using the structured clinical interview for the International Statistical Classification of Diseases and Related Health Problems (ICD-11)[49]. Patients included in the study were euthymic for at least 2.5 months before recruitment. Additional inclusion criteria were: most recent episode being depression, aged 18-50 years, absence of symptoms from other mental health conditions beyond BD, and no history of substance abuse. We assessed residual mood symptoms and cognitive performance using validated scales on mania, anxiety, depression and tasks on executive and general cognitive performance. See **Table 1** for further details, and **Supplemental Material** for sample size estimates (minimum sample size estimated was 20 per group, to identify volatility effects at 0.8 power). The study was approved by the Institutional Review Board of National Research University Higher School of Economics and the Local Ethical Committee of the First Moscow State Medical University. All participants provided written informed consent.

**Table 1.** Group demographics and variables describing the measures of affective state, cognitive and executive function in euthymic bipolar disorder patients (BD, N = 22) and healthy control participants (HC, N = 27). For MEG analysis, data were available for 27 HCs and 21 BDs. Values are provided as mean and SEM (M, SEM) or as count and percentage (n, %). Between-group differences in scale values were assessed using permutation tests. Our alpha significance level was set at 0.05, and we controlled the false discovery rate (FDR) at q = 0.05 for multiple tests. Permutation test p-values were complemented with Bayes factor analysis to assess the amount of evidence in favour of H_1_ or H_0_. Age and sex distribution were comparable across groups (Age: *P_FDR_*= 0.4921, no-significant differences; *BF_10_* = 0.3472, providing anecdotal evidence for H_0_; Sex: Chi-squared statistic, χ^2^*=* 1.6559, *df = 1, P* = 0.1849, no significant; *BF_10_* = 1.102, no evidence for H_0_ or H_1_). There was substantial and anecdotal evidence supporting no differences between BD and HC groups in anxiety, depression, mania, and general cognitive scores (Bayes factor, BF_10_, in the range 1/3–1: anecdotal evidence; 1/10–1/3: substantial evidence). Between-group differences after FDR control were observed exclusively for Part 2 of the Trail Making Test (denoted by *), assessing task switching, based on strong evidence. There was anecdotal evidence for differences in Wisconsin Card Sorting Test scores, assessing executive functions. However, this effect was not significant after FDR control. ^1^State anxiety scores on the day of the experimental session were available for 22 bipolar patients and 18 healthy participants. Abbreviations: HADS, Hospital Anxiety and Depression Scale. The type of mood stabiliser taken by bipolar participants was mainly Lamotrigine (n = 11), but a few patients were taking Carbamazepine (n = 2), Valproic acid (n = 2), or Lithium (n = 1). See **Supplementary Materials** for further details on the scales and the corresponding references.

|  | Bipolar group<br>(N = 22) | Healthy control group<br>(N = 27) | Statistics (Permutation test<br>p-value and Bayes factor) |
| --- | --- | --- | --- |
| Age (years), M (SEM) | 29.1 (SEM 1.67) | 27.5 (SEM 1.18) | $P = 0.4921$, $BF_{10} = 0.3472$ |
| Sex (females), n (%) | 17 (77.2%) | 15 (56%) | $\chi^2 = 1.6559$, $df = 1$, $P = 0.1849$; $BF_{10} = 1.102$ |
| Duration of illness (years), M (SEM) | 7.4 (SEM 1.04) | NA | NA |
| Last episode polarity, (%) | depression (100%) | NA | NA |
| Time in euthymia (months), M (SEM) | 4.9 (0.68) | NA | NA |
| Bipolar II, n (%) | 17 (77.3%) | NA | NA |
| Bipolar I, n (%) | 5 (22.7 %) | NA | NA |
| Antidepressant, n (%) | 10 (45.5 %) | NA | NA |
| Antipsychotic, n (%) | 14 (63.6 %) | NA | NA |
| Mood stabiliser, n (%) | 16 (81.8 %) | NA | NA |
| Summary: Taking psychiatric<br>medication, n (%) | 19 (86.4%) | NA | NA |
| HADS anxiety, M (SEM) | 5.5 (0.94) | 5 (0.64) | $P_{FDR} = 0.64$, $BF_{10} = 0.31$ |
| HADS depression, M (SEM) | 4 (0.69) | 3.30 (0.57) | $P_{FDR} = 0.4003$, $BF_{10} = 0.3824$ |
| Beck Depression Inventory, M (SEM) | 8.3 (1.75) | 5 (1) | $P_{FDR} = 0.1038$, $BF_{10} = 0.8668$ |
| Altman's Mania Scale, M (SEM) | 1.7 (0.50) | 2.6 (0.50) | $P_{FDR} = 0.2040$, $BF_{10} = 0.5652$ |
| State-Trait Anxiety Inventory (state<br>subscale on MEG day)<sup>1</sup>, M (SEM) | 35.5 (1.72) | 33.7 (1.90) | $P_{FDR} = 0.4571$, $BF_{10} = 0.3833$ |
| Trail Making Test (Part 1), M (SEM) | 23.3 (1.21) | 21.2 (1.22) | $P_{FDR} = 0.2216$, $BF_{10} = 0.5296$ |
| Trail Making Test (Part 2), M (SEM) | 87.8 (8.5) | 55.7 (5.6) | **$P_{FDR} = 0.0022^*$, $BF_{10} = 14.907$** |
| Wisconsin Card Sorting Test, M (SEM) | 29.3 (0.92) | 31.6 (0.67) | $P_{FDR} = 0.0364$, $BF_{10} = 1.6560$ |
| Mini-Mental State Examination, M<br>(SEM) | 29.5 (0.18) | 29.5 (0.17) | $P_{FDR} = 0.8678$, $BF_{10} = 0.2875$ |

### Reward-based motor sequence learning task

Participants underwent an initial fine motor control assessment, then completed a validated motor-based decision-making paradigm[48] (**Figure 1A**), which combines probabilistic binary reward-based learning within a volatile setting (reminiscent of reversal learning) with the execution of motor sequences to express decisions. Participants learned two sequences of four finger presses, followed by a 320-trial test phase. In each trial, they were required to choose and perform one of the sequences to potentially earn a reward (5 points; **Figure 1A**). Reward probabilities for sequences were reciprocal (p, 1-p) and changed pseudorandomly (**Figure 1B**). The aim was to infer the reward probability associated with each sequence (’action-outcome’ contingencies henceforth) and adjust their choices considering changing contingencies. Accumulated points translated to monetary rewards. See timeline in **Figure 1C**. The task, programmed in MATLAB using Psychtoolbox, recorded participants’ keypress timings to evaluate reaction time (RT) and performance tempo (**Figure 1D**). See **Supplemental Materia**l.

**Figure 1.**
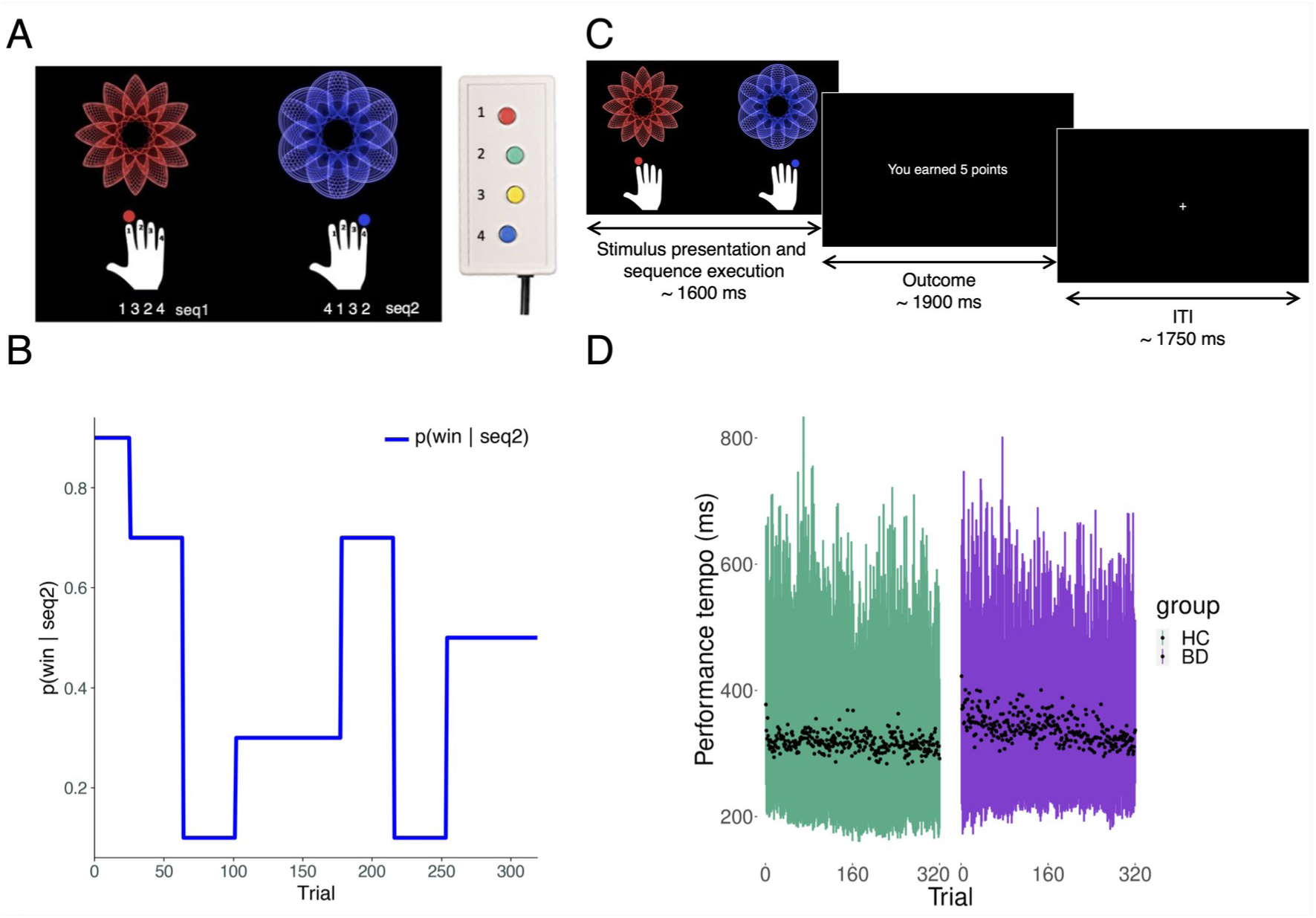
Experimental paradigm and motor performance overview. a. The initial phase of the task involved practicing two motor sequences, each linked to a distinct fractal image. The red fractal corresponded to sequence 1 (seq1: 1-3-2-4), and the blue fractal to sequence 2 (seq2: 4-1-3-2), with button presses producing sounds of varying pitches (E5, F5, G5, A5). **b.** The stimulus-outcome mapping varied per participant across each block of 160 trials, with the win probability shifting every 26-38 trials through different phases (blue fractal: *p(win|seq2)* = 0.9, 0.7, 0.1, 0.3, 0.5) and the red fractal (seq1) having reciprocal probabilities (*p(win|seq1) =* 1-*p(win|seq2)*). **c.** Each trial presented the fractals on-screen, prompting participants to perform the sequence they believed most likely to win, aiming to maximize rewards. Binary feedback on reward acquisition was displayed 1250 [± 250] ms after sequence performance, visible for 1900 [± 100] ms, indicating either ‘You earned 5 points’ or ‘You earned 0 points’. **d.** Displays the trial-by-trial performance tempo (ms) for the healthy control (HC, green) and bipolar disorder (BD, purple) groups. Tempo, calculated as the mean inter-key press interval, is shown as trial-wise averages (black dots) with 95% confidence intervals represented by bars.

### MEG recording and preprocessing

MEG was performed using a 306-channel system (Elekta Neuromag VectorView), with head movements tracked by a head position indicator with four coils. Concurrently, ECG and EOG were recorded for MEG artefact rejection. Recordings were sampled at 1000 Hz and filtered between 0.1–330 Hz. MEG preprocessing involved head movement correction, noise reduction, and channel selection using standard methods ([50]; Elekta Maxfilter software; **Supplemental Material**). MEG data was further processed using MNE-python[51] (Python version 3.11.5) and custom Python scripts, lowpass filtered at 125 Hz, downsampled to 250 Hz, with a notch filter applied at 50 and 100 Hz. Independent component analysis (FastICA) removed eye and heart artifacts (3.3. ICs on average per participant).

### General task performance

General task performance was assessed using the win rate (rewarded trials), average tempo (mean inter-key press interval in a sequence, mIKI), and RT (time from stimulus to first key press).

To assess practice effects in performance tempo and RT over trials as a function of the group, we employed Bayesian multilevel regression modelling (BML) with the logarithm of trial-wise RT or mIKI (in milliseconds) as the dependent variables. BML was performed with the brms R package[52,53]. For each dependent variable, models of decreasing complexity were constructed (**Table S1**). Details on model comparison can be found in the **Supplemental Material**.

### Modelling decision-making behaviour using hierarchical Gaussian filters

To assess probabilistic learning in our task we used a validated hierarchical Bayesian model, the 3-level perceptual HGF for binary categorical inputs[23,24](**Figure 2a**). This model described how participants infer hidden states about the tendency of the action-outcome contingencies on trial *k*, *x_2_ ^(k)^*(level 2), and the rate of change in that tendency (log-volatility), *x_3_ ^(k)^*. Level 1 represents the binary reward input. Gaussian belief distributions on levels 2 and 3 are represented by their posterior mean (*μ_2_ ^(k)^*, *μ_3_ ^(k)^*) and posterior variance (uncertainty: *σ_2_*, *σ*_3_). Precision is the inverse variance or uncertainty, *π_i_ (i* = 2, 3). Belief updating on each level *i* and trial *k* is driven by prediction errors, and modulated by precision ratios, weighting the influence of precision or uncertainty in the current level and the level below. This is termed precision-weighted PE, pwPEs. For level 2, belief updating takes the simple form:

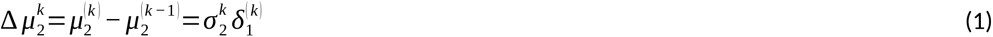

**Figure 2.**
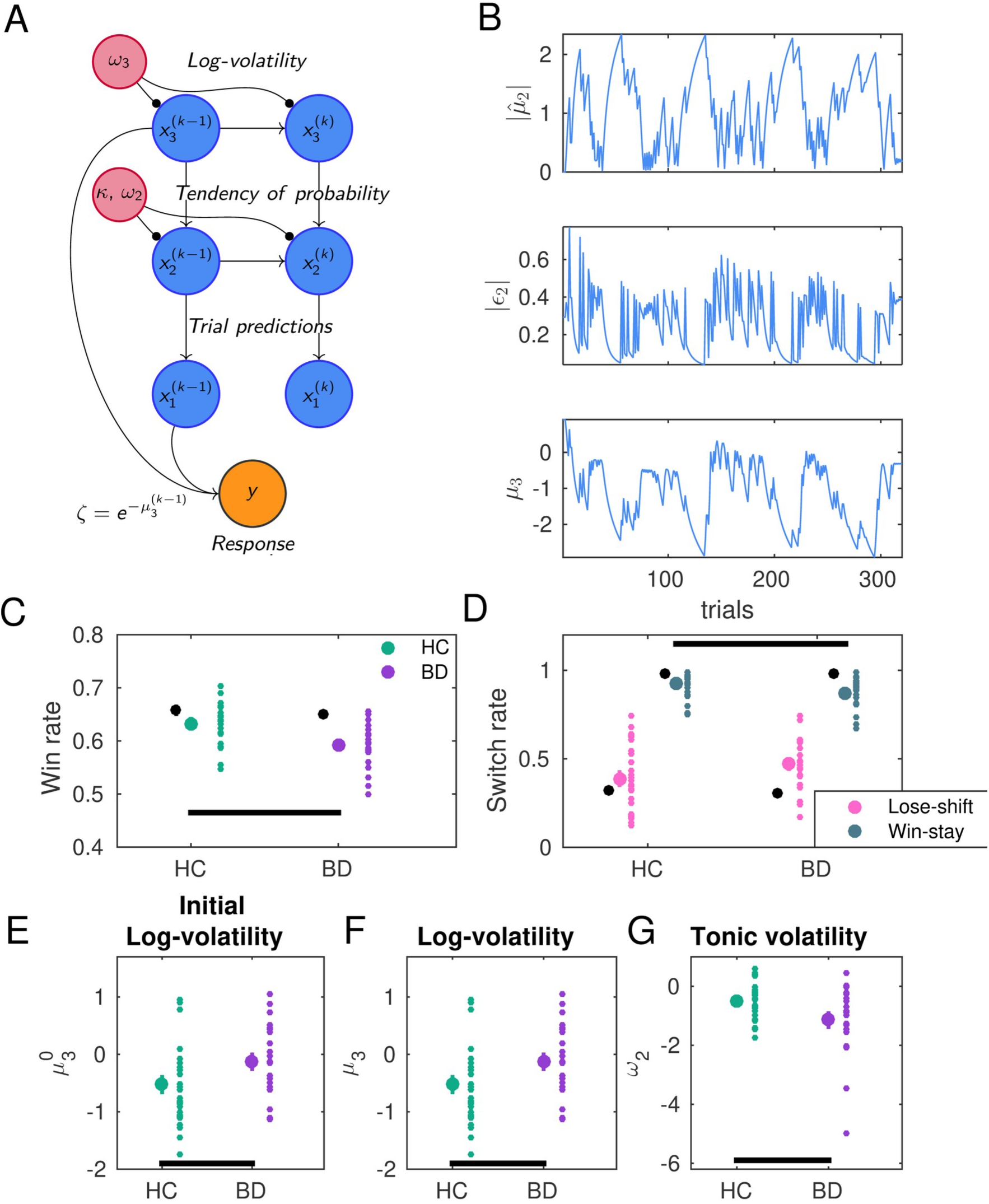
Computational model and changes in learning behaviour in euthymic bipolar patients. a. Overview of the winning model: 3-level binary categorical HGF perceptual model and coupled response model. In this model, agents infer the current tendency of the action-outcome probabilistic mapping on trial *k*, *x_2_^(k)^*, and its rate change or log-volatility, *x_3_^(k)^*. Beliefs about these true states are updated using one-step equations parametrised by their mean (*μ_2_^(k)^*, *μ_3_^(k)^*) and variance (*σ_2_^(k)^*, *σ*_3_*^(k)^*), representing uncertainty or the inverse of precision). These updates are modulated by parameters, such as *κ, ω*_2_, *ω*_3_. The response model maps these beliefs to decisions based on the prediction of log-volatility from the previous trial (*μ_3_^(k-1)^*). **b.** Trajectories used in further analyses include the strength of predictions about action-outcome contingencies,|*μ̂*_2_^(*k*)^| (top), for assessing motor vigour effects; the trajectory of unsigned precision-weighted prediction errors updating beliefs at level 2, labelled|ε_2_| here (centre), serving as a parametric regressor of source-reconstructed MEG activity, alongside uncertainty regressors *σ_2_*, *σ*_3_; and log-volatility estimates, *μ_3_* (bottom), averaged to test the hypothesis that BD participants overestimate volatility in this setting. **c.** Comparative win rates show BD participants (purple) were significantly less successful in achieving rewarding outcomes than their healthy counterparts (green; lower win rate, *P*_FDR_ = 0.0014, permutation test). **d.** BD patients exhibited a significantly higher tendency to switch after a win compared to the HC group (reduced win-stay behavior, *P*_FDR_ = 0.0194). Nonetheless, lose-switch behavior was similar across groups (*P* = 0.0966, non-significant; BF_10_ = 0.8905; anecdotal evidence against group differences). Mean and SEM rates shown in black dots for panels c and d represent performance by ideal Bayesian observers with the same input as our participants (detailed in **Supplemental Material**), highlighting deviations from these ideal patterns in our actual participants, which however did not account for the observed between-group differences. **e-g.** Between-group comparisons of HGF computational variables revealed that BD patients consistently overestimated environmental log-volatility (**e**; initially, *μ_3_*^(0)^: *P*_FDR_ = 0.0142, and throughout the task, **f**; mean *μ_3_*: *P*_FDR_ = 0.0428), while showing an attenuation effect on tonic volatility, *ω_2_* (**g**; significant reduction compared to HC, *P*_FDR_ = 0.0174).

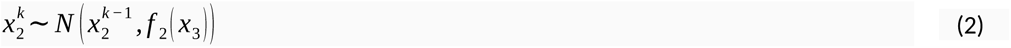

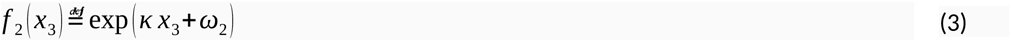

In (3), *ω*_2_ represents the invariant (tonic) portion of the log conditional variance of *x*_2_, while *κ* is a coupling constant that modulates the influence of phasic volatility, *x*_3_, on the size of action-outcome belief updates. The step size on level 3 is modulated by *ω*_3_, a high-level tonic volatility parameter. See **Supplemental Material** for modelling details.

To assess how beliefs mapped to decisions, we coupled this perceptual model to response models previously used in similar tasks[24,37,38]. First, we considered a unit-square sigmoid response model where choice probability is shaped by a free fixed (time-invariant) parameter ζ, interpreted as inverse decision noise: the sigmoid approaches a step function as ζ tends to infinity. This constituted our model M1. Model M2 was similar but employed a two-level HGF with constant volatility. M3 combined the 3-level HGF with a response model where the sigmoid function depends on the trial-wise prediction of log-volatility, 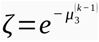 [22](**Figure 2a**). In this model, higher estimates of volatility lead to a more stochastic mapping from beliefs to decisions. As a result, there is an increased likelihood of choosing responses that deviate from predictions, consistent with increased exploration (exploring whether the contingency has changed). In models M1 and M3, parameters *ω*_2_ and *ω*_3_ were free; *ω*_2_ was also free in M2. Additionally, ζ was free in M1 and M2, while initial values *μ_3_^(0)^* and *σ_3_^(0)^* were free in M3. A fourth model, M4, was constructed similarly to M3 but replaced the free parameter *ω*_2_ with κ[28].

Models were fit to individual behavioural data (series of chosen and rewarded sequences: responses and observed outcomes), using prior values described in **Table S2**. Log model evidence was used for model comparison using random-effects Bayesian model selection (**Supplemental Materials**). As in previous work, simulations were conducted to quantify how well free model parameters could be estimated[27,37]. Relevant belief and uncertainty trajectories were used subsequently for our MEG analysis (**Figure 2b**; see **Results**). The models were implemented as a part of the TAPAS toolbox[54]. We used the HGF release v7.1 in Matlab R2020b, and functions ‘tapas_ehgf_binary’.

### Assessing motor invigoration

We additionally investigated whether trial-by-trial predictions about the probabilistic action-outcome mapping, *μ̂*_2_^(*k*)^, differentially influenced motor performance in our groups. Following Tecilla et al.[48], given the sign of *μ̂*_2_^(*k*)^ is arbitrary, we used as predictor for Bayesian multilevel modelling the absolute value,|*μ̂*_2_^(*k*)^|, representing the *strength* of those predictions, and assessed its association with performance variables (log-mIKI, log-RT). We hypothesised a negative correlation, suggesting that stronger expectation about reward contingencies speeds performance. We also hypothesised a greater sensitivity to these predictions (steeper slope) in BD compared to HC. Details on models are in **Table S3**.

### Source reconstruction of MEG signals

We reconstructed MEG signals using Linearly Constrained Minimum Variance beamforming (LCMV[55]) in MNE-Python and individual T1-MRI images for cortical divisions with Freesurfer 6.0[56,57], http://surfer.nmr.mgh.harvard.edu/). We aligned MRI and MEG coordinate systems, selected the Desikan-Killiany–Tourville atlas for cortical parcellation (DKT[58]), and performed forward modelling with boundary element models[37].

We focused on alpha, and beta frequency bands, band-pass filtering signals between 1–40 Hz before LCMV beamforming. Theta-band activity was also examined given its robust association with feedback processing[34,59], relevant for win/lose outcomes in our task. Gamma frequency analysis followed a similar process (30–124 Hz band-pass filter). Time courses were extracted for regions of interest (ROIs) associated with decision-making under uncertainty and reward processing[35,60–65], and linked with impairments in fronto-striatal reward circuitry in [5,10,66,67]. These included the (1) ACC, (2) OFC, including the ventromedial PFC, (3) dorsomedial PFC (dmPFC), (4) dorsolateral PFC (dlPFC). We also included the (5) primary motor cortex (M1) and (6) premotor cortex (PMC), to assess motor activity during decision-making[68].

Our study’s ROIs comprised 16 bilateral labels in eight areas from the DKT atlas: (1) rostral and caudal ACC, (2) lateral and medial OFC (including vmPFC), (3) superior frontal gyrus (dmPFC, and supplementary motor area, SMA), (4) rostral middle frontal gyrus (rMFG), (5) precentral gyrus (M1), and (6) caudal MFG. Time series extraction utilised the PCA flip method in MNE-Python. Although the ’flip’ operator was not relevant for our time-frequency analysis, it was essential for preparing the source-reconstructed time series for subsequent connectivity analysis. See anatomical label references in **Supplemental Material**.

### Convolution modelling of time-frequency responses during outcome processing

We used a validated convolution-modelling approach to analyse frequency-domain amplitude changes related to belief updating and uncertainty following outcome presentation[36,38,69]. Building on previous work[37], this frequency-domain general linear model (GLM) included as parametric regressors the unsigned pwPE updating beliefs on level 2 (representing precision-weighted Bayesian surprise; the absolute value is preferred for the binary HGF where sign on level 2 is arbitrary[70,71]), and uncertainty measures (σ_2_, σ_3_). It also included discrete regressors for win/lose outcomes and error trials. To avoid regressor collinearity and potential GLM misspecification, we excluded the level 3 pwPE[37,38], due to its high linear correlation with the unsigned pwPE on level 2 (**Supplementary Materials**).

The GLM was applied to concatenated epochs of source-reconstructed data in our ROIs, using Morlet wavelets for time-frequency (TF) analysis in 4–100Hz and within −0.5–1.8 s (**Figure S1**). We hypothesised between-group differences in gamma and concomitant alpha/beta activity during pwPE processing, and in alpha/beta activity during uncertainty encoding. Theta 4–7 Hz activity was assessed for the win/lose regressors[34,59].

We conducted this analysis using SPM12 software (http://www.fil.ion.ucl.ac.uk/spm/), adapting original code by Spitzer et al.[72], as used in Hein et al.[37], with additional details available in the **Supplemental Materials**.

### Frequency-resolved functional connectivity

To analyse directed functional connectivity between frequency-resolved activity in our ROIs, we employed time-reversed Granger causality (TRGC[73]; **Supplemental Materials**) as a robust metric for directed information flow[74]. Following the recommendations of Pellegrini et al.[74], we applied TRGC in the frequency domain to LCMV-based source-reconstructed time series from our 16 ROIs after the PCA flip transformation.

Our analysis focused on between-group differences in the directionality of information flow within the 8–30 Hz range during the 0.5–1 s interval of outcome processing for trials with large unsigned pwPEs updating beliefs at level 2. We employed a median split of unsigned pwPE values, yielding approximately 160 high-| pwPE| trials per participant. This frequency range was selected based on evidence that beta-band functional connectivity from the PFC effectively differentiates levels of predictability, exhibiting reduced values during unpredictable trials[34]. By examining TRGC in trials with high unsigned pwPE values, we anticipated a general decrease in beta-band TRGC in HC, in parallel with alpha/beta amplitude suppression during belief updating. We hypothesised that this pattern would be disrupted in BD. The TRGC analysis was conducted using the ROIconnect plugin for EEGLAB[74], adapted for our MNE-python LCMV outputs. See **Supplemental Materials**.

### Statistical analysis

Between-group analyses of behavioural, computational, and TRGC-derived variables used independent-sample permutation tests (5000 permutations) in MATLAB®. Within-subject analyses used paired permutation tests. We maintained an alpha significance level at 0.05 and controlled false discovery rates (FDR) at *q* = 0.05 for multiple tests. Non-parametric effect sizes are reported as probability of superiority[75,76] (Δ). Non-significant effects were further evaluated using Bayes Factors (BF _10_), interpreted following Wetzels and Wagenmakers[77].

Statistical analysis of source-level time-frequency images used cluster-based permutation testing in the FieldTrip Toolbox[78,79] (1000 permutations). We averaged TF activity across frequency bins within each band (theta, alpha, beta; 60–100 Hz for gamma[37]). Temporal intervals of interest for statistical analyses were selected based on previous research[37,38,59]: 0.5–1.8 s for parametric regressors, 0.2–1 s for win/lose regressors. We controlled the family-wise error rate (FWER) at 0.025. See **Supplemental Materials**.

## Results

### Demographics

BF analysis provided anecdotal evidence for a balanced distribution of age and sex across the groups. Furthermore, substantial to anecdotal evidence indicated similar scores in mania, anxiety, depression, and general cognitive functioning between groups. Significant differences were observed exclusively in executive functioning, with the BD group demonstrating lower performance. See **Table 1**.

### Altered reward-based decision dynamics in bipolar disorder during euthymia

Euthymic BD patients exhibited lower win rates compared to HC individuals (*P*_FDR_ = 0.0014; Δ = 0.79, CI = [0.60, 0.90]; **Figure 2c**). They also demonstrated lower win-stay rates (*P*_FDR_ = 0.0194; Δ = 0.71, CI = [0.55, 0.85]; **Figure 2d**). This indicates that, after securing a win on a trial, BD individuals were less likely to repeat the sequence compared to HC. Their decision to switch strategies post-loss was similar, based on anecdotal evidence (lose-shift rate: *P* = 0.0966; *BF_10_* = 0.8905; **Figure 2d**), and despite an overall increased total switch rate in BD relative to HC (See details in **Supplemental Materials**, including evidence for similar performance error rates).

To test our computational hypotheses, we used the HGF framework[24]. Bayesian model selection identified as the best model overall, and for each group separately, a three-level HGF with a response model in which the decisions depend on dynamic trial-by-trial expectations of log-volatility, *μ_3_^(k-1)^*, and with *ω*_2_, *ω*_3_, *μ_3_^(0)^*, and *σ_3_^(0)^* as free model parameters (M3; **Table S4**). Simulation analyses confirmed good parameter recovery (**Figure S2**).

Using this model, we found that BD participants had higher expectations of log-volatility initially and on average (*μ_3_^(0)^*: *P*_FDR_ = 0.0142; Δ = 0.70, CI = [0.55, 0.85]; trial-average *μ* : *P* = 0.0428; Δ = 0.66, *CI* = [0.52, 0.82]; **Figure 2ef**). This suggests increased stochasticity in their responses, as also indicated by a correlation between log-volatility *μ_3_* and the response switch rate (**Figure S3**). Additionally, BD participants exhibited lower tonic volatility, *ω_2_*, compared to HCs (*P_FDR_*= 0.0174; Δ = 0.67, CI = [0.52, 0.82]; **Figure 2g**), suggesting a slower adjustment of beliefs about action-outcome contingencies (see simulation analysis in **Figure S4**). No significant between-group differences were found in *ω_3_*.

We additionally assessed the association between residual symptoms in BD participants and relevant HGF variables. Prior work suggests a positive correlation between volatility and trait anxiety[37]. Accordingly, we analysed the relationship between trait anxiety levels in BD and *μ_3_*, confirming significant positive correlation (Spearman’s rank correlation *ρ* = 0.46, 95% confidence interval [0.04, 0.75], *P_FDR_* = 0.030). For mania scores, we hypothesised a correlation with the precision weights term, *σ_2_*, which scales the influence of prediction errors on belief updates about action-outcome contingencies (eq.1). The rationale was that higher mania levels in BD might be associated with an enhanced reactivity to PEs[19], leading to faster belief updating driven by *σ_2_*. Linear regression analyses revealed a negative association between mania and *σ_2_* (*ρ* = -0.46 [-0.75, -0.02], *P_FDR_* = 0.037). Conversely, we considered that depression scores might be associated with attenuated reward-based belief updating (lower *σ_2_*), yet found a lack of association between these variables (*ρ* = 0.04 [-0.34, 0.41], *P* = 0.836; *BF_10_* = 0.464, anecdotal evidence). See **Figure S5** and **Supplemental Materials**.

### Similar timing of actions during motor decision-making in euthymic bipolar and healthy participants

Bipolar participants, although slower in baseline motor performance (**Supplemental Materials**), matched HC in tempo during the main task (mIKI: HC 339 [16.6] ms, BD 354 [18.2] ms, *P* = 0.6223; *BF_10_* = 0.33, substantial evidence against H_1_). RTs were also comparable between groups (meanRT HC 501 [20.1] ms, BD 553 [32.2] ms, *P* = 0.2262; *BF_10_* = 0.55).

Additionally, both groups improved in tempo and RT over trials, as expected, but BD participants exhibited faster improvements in tempo (**Figure 3a-c)**. Bayesian multilevel modelling confirmed this (**Table S5; Figure S6)**. The best model demonstrated a negative slope in practice effects for both timing variables (negative posterior point estimate, with the 95% credible interval [CI] not including zero, denoting a credible non-zero Bayesian estimate; **Figure 3ab; 3d-f**). The slope of the practice effect on tempo was steeper in BD (95% CI with negative slope difference; **Figure 3c**). See **Supplemental Materials**.

**Figure 3.**
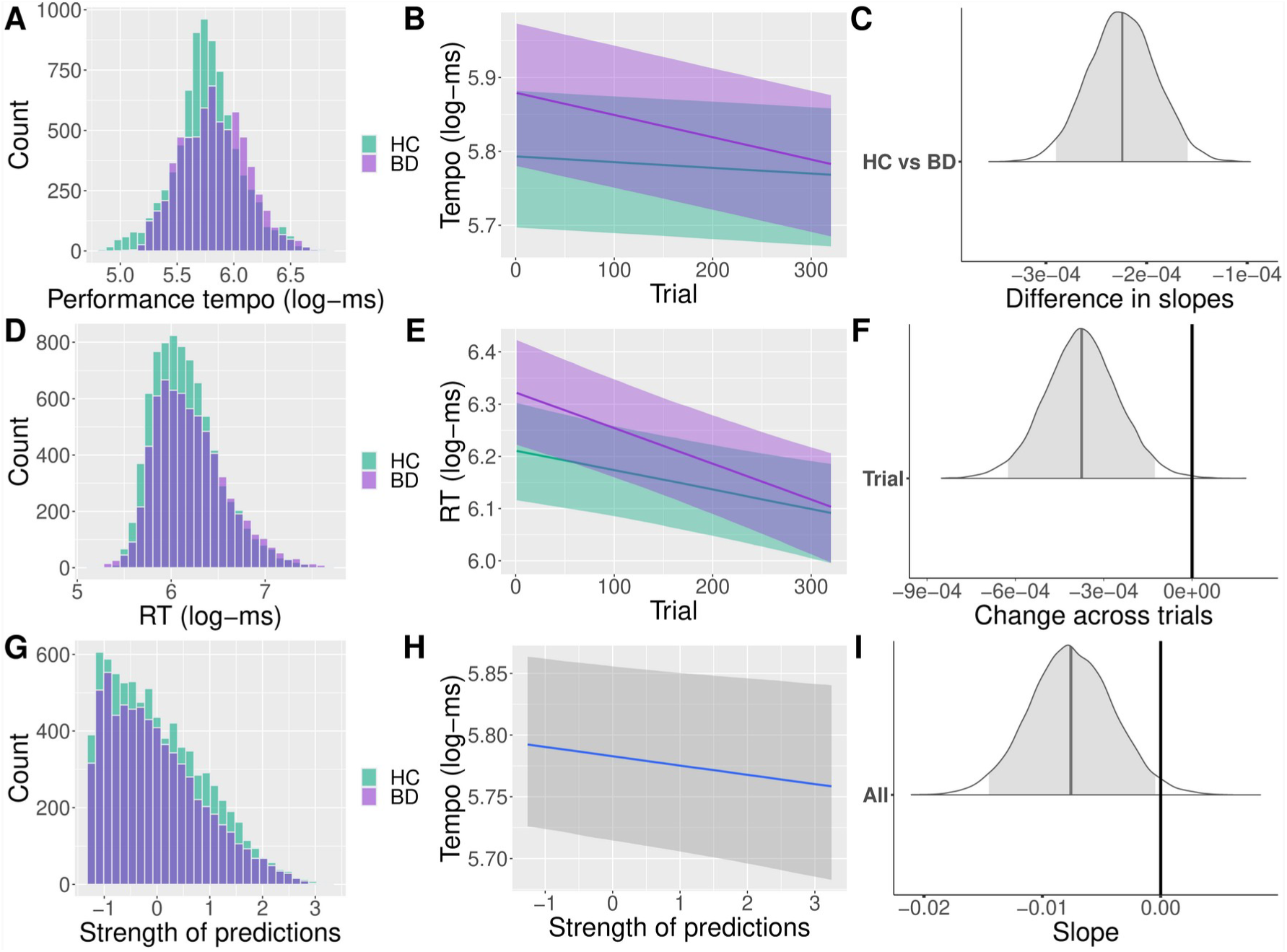
Effects of practice and expectations of action-outcome contingencies on performance timing. **a-c.** Practice effects on performance tempo, measured as the logarithm of the mean inter-key press interval (log-mIKI, in log-ms). a. Distribution of log-mIKI (“tempo” in the graphics) in the BD (purple) and HC (green) cohorts. **b**. Output of the Bayesian multilevel model that provided a better fit to the data (see main text). The graphic represents the association between trial number and performance tempo, represented as log-mIKI, in each cohort. The bold lines and shaded areas denote the posterior mean and the 95% credible interval (CI, shaded areas) in each group. In the reference HC group, participants sped up their performance across trials. **c.** Illustration of the posterior distribution of the between-group difference in slopes, including the posterior point estimate (grey vertical line) and 95%-CI (denoted by the grey area under the curve). The BD minus HC slope difference was negative, and the 95%-CI did not overlap with zero, supporting that performance in the BD group sped up more throughout the task. **d-e**. Same as a-c but for the logarithm of reaction time, log-RT (in log-ms). The reference HC group also demonstrated a reduction in RT over trials (panel e). The 95% CI of the between-group slope difference included zero, indicating a credible effect for a similar change over time in RT in both groups. **f.** Posterior distribution of the fixed effect of trials, showing negative changes in RT over time. The black vertical line represents a zero change difference, which did not overlap with the 95% CI of the estimate, suggesting a 95% probability of a negative change in RT over trials (speeding up). **g-i.** Invigoration of performance tempo by the strength of predictions about the tendency of the action-outcome contingencies. **g**. Histogram of the strength of predictions about reward contingencies |*μ̂*_2_^(*k*)^| in BD and HC groups, showing similar distributions. **h**. Bayesian multilevel modelling demonstrated that performance tempo (log-mIKI, in log-ms) was shorter in trials when participants had a stronger expectation on the reward tendency, as the posterior estimate of the slope of this association was not negative, and the 95%-CI did not include zero (**i**). Thus, in the joint cohort, individuals speed up sequence performance when they hold stronger predictions about the action-outcome contingencies.

### Expectation about reward probability invigorates motor performance similarly in euthymic bipolar and healthy participants

Further analysis revealed that expectations about the tendency of the action-outcome probability similarly influenced performance tempo in BD and HC groups. BML identified as the best model one in which performance tempo (log-mIKI) was modulated by variations in the predictor|*μ̂*_2_^(*k*)^|, without necessitating the inclusion of group differences (**Supplemental Materials; Table S6**). This model demonstrated that stronger predictions led to faster performance across all participants, irrespective of group, confirmed by the negative slope of this association (95% CI not including zero; **Figure 3gi; Figure S7**). Conversely, log-RT was not modulated by the strength of predictions or other variables (**Table S6)**.

### Attenuated neural representation of precision-weighted prediction errors updating beliefs about the action-outcome contingencies in bipolar disorder

During the processing of unsigned pwPEs about the tendency of action-outcome contingencies, HC and BD participants exhibited suppression of 8–30 Hz activity across prefrontal, orbitofrontal, cingulate, and motor regions (negative cluster within 0.5–0.9 s, post relative to pre-outcome baseline, *P_FWER_* = 0.001, 0.024 in each group; **Figure 4ab)**. This suppression effect was less widespread in BD, and the between-group difference was significant across the caudal and rostral ACC, MFG, and OFC; as well as in the SFG and M1 (BD – HC positive cluster at 0.6–0.9 s, *P_FWER_* = 0.0130; **Figure 4cd; Figure S8**). Alongside these 8–30 Hz effects, the BD group exhibited significantly attenuated high gamma activity (60–100 Hz) compared to HC (negative cluster, *P_FWER_* = 0.0090; **Figure 4e**). The latency of the gamma effect coincided with the timing of the alpha-beta modulations, spanning 0.5–0.82 seconds, and overlapping within the aforementioned ROIs. No significant within-subject changes in gamma activity to the unsigned pwPE regressor were observed in either group (**Supplemental Material**).

**Figure 4.**
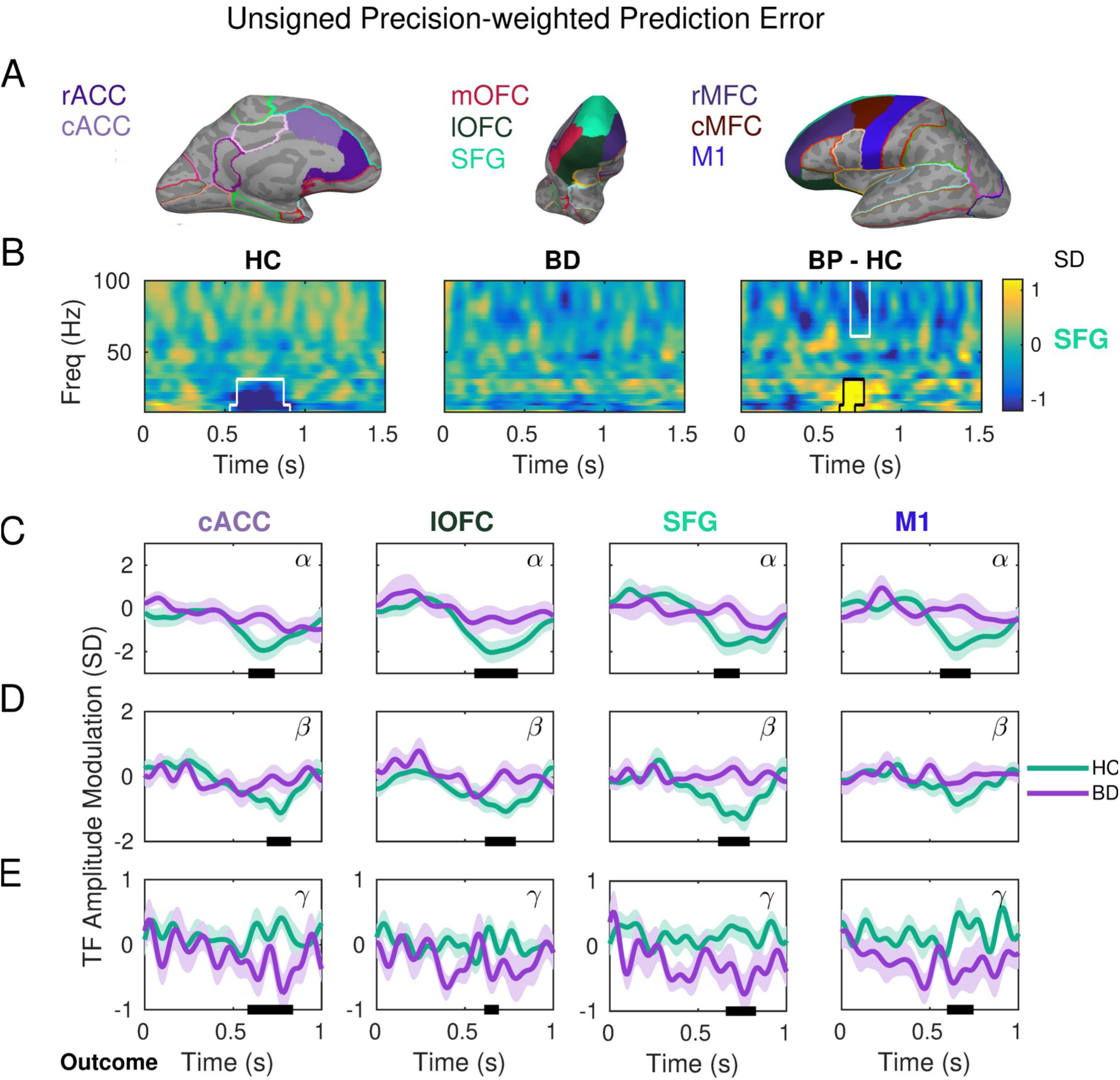
Attenuated gamma increase and alpha-beta suppression during encoding unsigned precision-weighted prediction errors about stimulus outcomes in bipolar disorder. a. Source reconstruction of MEG signals was carried out with linearly constrained minimum norm variance (LCMV) beamforming. The statistical analysis of convolution GLM results targeted brain regions implicated in decision-making under uncertainty and reward processing[35,61–65], associated with impairments in the fronto-striatal reward circuitry in BD[5,10,66,67]: caudal and rostral ACC, OFC (lateral and medial portions: lOFC, mOFC), SFG, caudal and rostral MFG, M1. Panel a illustrates these regions using anatomical labels from the neuroanatomical Desikan-Killiany–Tourville atlas (DKT), utilised to parcellate the cerebral cortex of each participant based on their individual T1-weighted MRI. **b.** Left and centre panels display within-subject effects in time-frequency (TF) images representing oscillatory amplitude responses to unsigned precision-weighted PEs about stimulus outcomes. TF images cover the 4–100 Hz range, including theta (4–6 Hz), alpha (8–12 Hz), beta (14–30 Hz), and gamma (32–100 Hz) activity. The TF images were normalised by subtracting the mean and dividing by standard deviation (SD) of the activity in the [−300, −50] ms pre-outcome interval, and thus are presented in SD units. Significant within-subject effects are outlined in black for the HC (left) and BD (centre) groups (cluster-based permutation tests, negative cluster within 0.5–0.9 s post relative to pre-outcome baseline, *P_FWER_* = 0.001, 0.024 in each group, respectively). The right panel shows the between-group differences, significant in a cluster-based permutation test (positive cluster within 8–30 Hz, *P_FWER_* = 0.0130; negative cluster within 60–100 Hz, *P_FWER_* = 0.0090; N = 21 BD and 27 HC independent samples). The time point 0 s marks the onset of outcome presentation. **c,d** Panels Illustrate between-group effects in the alpha (c) and beta (d) ranges, attributed to more pronounced alpha and beta suppression in HC than in BD participants during encoding of unsigned pwPE on level 2. Effects are depicted in ROIs including the cACC, lOFC, SFG, M1. **e**. Similar to panels c and d but in the gamma range, showing that unsigned pwPE were associated with increases in TF amplitude in gamma range for HC participants, yet with gamma attenuation in BD participants, and across a similar range of ROIs. Labels denote the rostral anterior cingulate cortex, rACC; caudal ACC, cACC; superior frontal gyrus, SFG; lateral and medial orbitofrontal cortex, lOFC and mOFC; primary motor cortex, M1; caudal and rostral middle frontal gyrus, cMFG, rMFC.

In addition, for the uncertainty regressors σ_2_ and σ_3_, despite a significant widespread increase in 8–30 Hz activity to estimation uncertainty σ_2_ in HC, no significant between-group differences were observed (**Figure S9**). Regarding theta modulation by win and lose events, no significant differences were observed between groups either. However, as expected, both groups showed significant increases in theta activity from baseline in the ACC, extending to prefrontal and orbitofrontal ROIs (**Figure S10**).

In a post-hoc analysis, we investigated alpha and beta raw power during inter-trial intervals. This aimed to determine whether BD’s reduced suppression in the 8-30 Hz range to the pwPE regressor indicated a limited dynamic range of activity at these frequencies. Significantly lower power was observed in BD compared to HC, yet exclusively at 13–20 Hz. This effect emerged in most of the ROIs where the pwPE effect was expressed (**Figure S11**; **Supplemental Material**).

### Frequency-domain functional connectivity patterns during unsigned pwPE processing

We next assessed group differences in the directionality of information flow during outcome processing for trials with large unsigned pwPEs updating beliefs at level 2. The BD cohort exhibited significantly larger TRGC coefficients than HC participants from the cACC to the rMFG and rACC, as well as from the SFG to the cMFG, in the beta frequency range (*P_FDR_* = 0.0032, 0.0064, 0.0064, respectively; **Figure 5**). The effect from the cACC to the rMFG extended to the alpha range (**Figure 5a,c**). These findings indicate stronger evidence for statistical dependencies between sources in the identified directions for BD than HC in the beta (alpha) range. Importantly, these between-group effects were not attributable to differences in signal-to-noise ratio (**Figure S12**).

**Figure 5.**
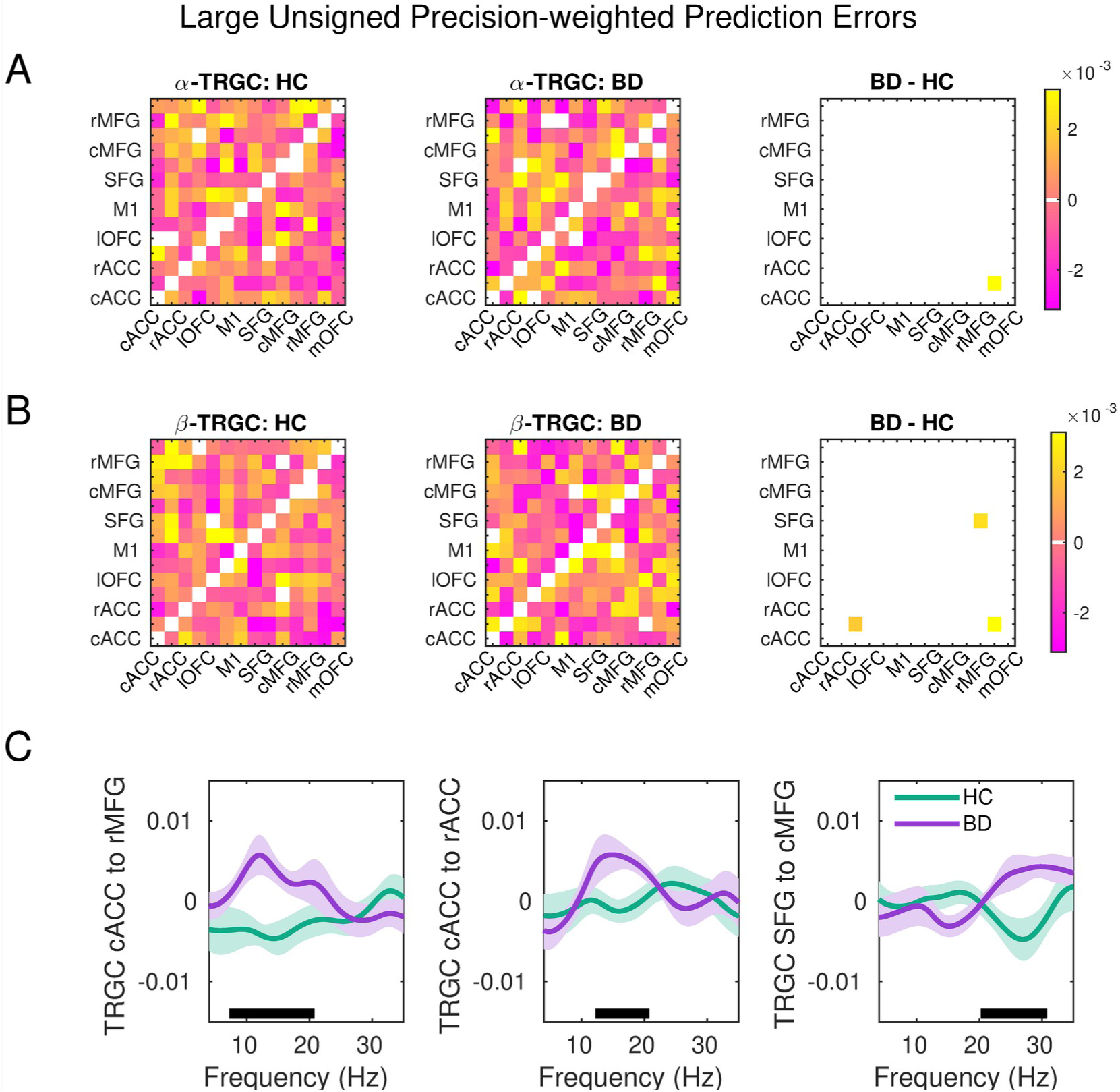
Time-reversed Granger causality during outcome processing for trials with large unsigned pwPEs updating beliefs at level 2. a. Comparison of TRGC estimates in the alpha band for healthy control participants (HC, left column), bipolar disorder patients (BD, centre), and their difference (BD-HC, right column). The direction of information flow goes from rows to columns, with positive coefficients denoting increased predictability in that direction, while negative coefficients denote the reverse (increased predictability from column to row). Between-group statistical analysis was conducted in the above-diagonal values. Anatomical labels represent our regions of interest, bilaterally. Labels are displayed for one hemisphere. The coloured pixel in the right panel indicates a significant between-group difference in TRGC metric, after FDR control, due to increased evidence for TRGC from the caudal ACC to the rostral MFG in BD (*P_FDR_* = 0.0032). **b.** Same as (a) but for the beta band, illustrating a significantly larger TRGC metric in BD than HC from the cACC to the rMFG and rACC, as well as from the SGF to the cMFG (*P_FDR_* = 0.0032, 0.0064, 0.0064, respectively). **c**. Left: Illustration of the TRGC metric from cACC to the rMFG between 8-30Hz for HC (green line: mean and SEM as shaded area) and BD (purple line: mean and SEM). The horizontal black line denotes the frequency bins of significant differences after FDR control, shown in (a). Middle: Same as the left panel but for the TRGC metric from cACC to rACC, showing beta effects. Right: Same as left and middle panels, exhibiting larger TRGC metric values in BD than HC from SGF to the cMFG in the beta range. Labels: rACC, rostral anterior cingulate cortex; cACC, caudal ACC; cMFG, caudal medial frontal gyrus, rMFG, rostral MFG; lOFC, lateral orbitofrontal cortex; mOFC, medial OFC; SFG, superior frontal gyrus; M1, primary motor cortex.

## Discussion

Completing our reward-based motor decision-making task, euthymic BD patients demonstrated lower win rates and a decreased tendency to repeat rewarded actions than HC, despite similar post-loss decision-making behaviour. Furthermore, employing the HGF to probe the computational processes underpinning decision-making, we found that BD participants expected more environmental volatility than HC, leading to a more stochastic mapping from beliefs to actions and higher switch rates, particularly after wins. These findings align with previous reports of heightened risk-taking and inconsistent behaviour in BD[80,81], mirroring elevated win-switch tendencies in BD adolescents[82] and deficits in response reversal during remission[3,4]. This suggests that decisions in euthymic BD are misaligned with their beliefs about recent successes, favouring suboptimal actions due to an overestimation of environmental changes, potentially overriding the influence of their beliefs about action-outcome contingencies on decisions.

Despite expecting increased volatility, BD patients were slower to adjust their expectations of action-outcome contingencies compared to HC, with a lower tonic volatility parameter ω_2_ indicating slower adaptation. Similar results in HGF modelling for paranoia[27] suggest that this propensity to anticipate change without learning from it appropriately may be a common feature across paranoia and bipolar disorder. Additionally, despite similar residual symptom levels in BD and HC, trait anxiety in BD correlated with volatility estimates, aligning with findings that high trait anxiety exacerbates difficulties in adapting to environmental changes[37,83].

Of particular relevance in BD, we observed that residual mania symptoms negatively correlated with estimation uncertainty, σ_2_, which scales the influence of PEs about action-outcomes on level-2 belief updating. Therefore, those with higher mania scores struggled more with updating these predictions. Given that early relapse in BD has been associated with a reduction in an empirical measure of belief updating in response to positive feedback[14], future work could investigate if computational metrics of belief updating like σ_2_ enhance prediction of clinical progression over behavioral indicators. Further investigations should also explore the effect of comorbid anxiety on volatility responses and relapse.

Despite deficits in decision-making and baseline executive function in our BD sample, motor performance invigoration was comparable to HC, indicating preserved motivational drive in euthymic BD. This contrasts with previous findings that rewards and success amplify energy and effort in BD[4,47]. Our Bayesian analyses revealed a similar sensitivity of performance tempo to expectations about reward contingencies in both groups, highlighting that the alterations in euthymic BD were confined to decision-making processes.

On a neural level, convolution modelling on source-reconstructed time-frequency activity revealed BD individuals had attenuated neural representations of encoding unsigned pwPE—updating beliefs about action-outcome contingencies—compared to HC. This was marked by decreased gamma and increased alpha-beta amplitude changes 0.5-0.9 seconds post-outcome across multiple PFC, OFC, ACC, and motor regions. Spatial effects in anatomical PFC labels corresponded with functional vmPFC, dmPFC, and dlPFC, aligning with the neural correlates of decision-making under uncertainty[35,62–64], and BD-specific neural alterations during reward processing[5,10,66,67].

Recent rhythm-based formulations of predictive coding suggest distinct roles of oscillatory activity at different frequencies in conveying predictions and PE during perception[33,84]. Alpha and beta oscillations in deep cortical layers are implicated in conveying top-down predictions, while gamma oscillations in superficial layers are associated with the representation of PE, particularly in sensory cortices and related areas[33,84]. This division has received empirical validation in both human and animal studies, supporting generative models like predictive coding[85–87] and hierarchical Bayesian inference[36,88], extending across perceptual and cognitive domains[35,89]. In models of hierarchical Bayesian inference, like the HGF, these oscillatory activities may underpin pwPE encoding, demonstrating an antithetical modulation of alpha/beta and gamma activity[36]. The observed dysregulation of these rhythms in conditions like anxiety[37,38] suggests a neurophysiological basis for symptoms resulting from imbalances in belief updating.

Our findings indicate that in euthymic BD, exacerbated alpha and beta activity may inhibit gamma activity during unsigned pwPE encoding, potentially accounting for maladaptive belief updating. This may reflect an under-reliance on using predictions about action-outcome contingencies to optimise behaviour, in line with the computational results. Such rhythmic changes match electrophysiological evidence of heightened beta and reduced gamma activity in BD during oddball processing[90,91]. Moreover, using the TRGC to assess directional influences in frequency-domain activity, we observed stronger evidence for beta-band directional flow in BD compared to HC, from cACC to rACC and rMFG and SFG to cMFG during trials with larger unsigned pwPEs. TRGC values increased in BD but decreased in HC, aligning with expectations from primate research where beta-band Granger Causality in the PFC decreases during unpredictable trials—a pattern suggesting normative responses[34]. Thus, our study revealed that euthymic BD was associated both with altered frequency-domain amplitude changes and functional connectivity during belief updating.

Insufficient GABAergic neurotransmission and excessive glutamatergic system activity have previously been linked to electrophysiological alterations in BD[92]. Considering that mood stabilisers for BD, such as valproate and lithium, may have opposing effects on beta activity and potentially on beta/gamma connectivity[92,93], a promising avenue for future research is to assess the modulation of alpha-beta and gamma amplitude and connectivity during pwPE encoding in BD as potential markers for tracking treatment response and for diagnostic purposes. A key limitation of our study is the inclusion of patients on diverse psychiatric medications, including mood stabilisers, antipsychotics, and antidepressants. These treatments impact various neurotransmitter systems like dopamine and serotonin, affecting neural and behavioural aspects of decision-making[9,94]. The varied effects of these medications may have influenced the magnitudes of the effects reported, a factor future research should consider. Additionally, the study was not preregistered, yet all analyses followed established pipelines from our recent work involving similar tasks[37,48], except for the TRGC analysis. This was specifically designed based on similar Granger-causality analyses that assess rhythm-based hypotheses of predictive processing[34]. Lastly, our study did not contrast the HGF framework with alternatives like Bayesian change-point models[95] or those jointly estimating volatility and stochasticity[21]. Future research would benefit from such comparisons to validate the computational processes underlying belief updating alterations in BD.

In sum, our findings highlight significant alterations in belief updating among BD individuals during euthymia, when learning reward-based probabilistic mappings in volatile environments, without affecting the motivational aspects of motor execution. Importantly, the identification of frequency-domain amplitude and functional connectivity alterations underpinning these computational maladaptations provides crucial insights for enhancing relapse prediction and monitoring treatment response in future research.

## Code availability

The code for the analyses will be shared at https://osf.io/m63a8/

## Conflict of Interest

The authors declare that they have no competing financial interests in relation to the work described.

## Funding

This research was partially supported by the Basic Research Programme of the National Research University Higher School of Economics (Russian Federation). The research used the Elekta Neuromag 306-channel MEG system at Centre for the neurocognitive research (MEG-Centre) in Moscow (Russian Federation).

## Notes

### Competing Interest Statement

The authors have declared no competing interest.

