## Supplementary Materials for "Frequency-specific changes in prefrontal activity associated with maladaptive belief updating in volatile environments in euthymic bipolar disorder"

### Supplemental Methods and Materials

#### Participants

The included bipolar patients met the criteria for euthymia for at least 2.5 months prior to recruitment, as indicated by the Beck Depression Inventory (BDI, [96]), Altman Self-Rating Mania Scale ([97]), Hospital Anxiety and Depression Scale (HADS, [98]), and State-Trait Anxiety Inventory (STAI, [99]). The healthy individuals did not have neurological or mental health conditions and were not currently taking any medication for anxiety or depression. Participants in both groups were right-handed, had normal or corrected-to-normal vision, and demonstrated the ability to perform controlled finger movements.

The BDI ranges from 0 to 63, with values above 14 denoting light to severe depression. The Altman Self-Rating Mania Scale ranges from 0 to 20, with values up to 5 considered normal. The HADS, which assesses anxiety and depression separately, ranges from 0 to 21, with normal scores in the range of 0 to 7. We additionally assessed state anxiety on the day of the experimental session to determine potential group differences, which could affect decision-making [70]. We used the state subscale of the Spielberger State-Trait Anxiety Inventory (Spielberger, 1983), ranging from 0 to 80, with values up to 30 considered low to medium.

In addition, in both cohorts, we assessed cognitive performance including executive functioning using the Trail Making Test (TMT, part 1 and 2, [100,101]), and reversal learning using the Wisconsin Card Sorting Test (WCST,[102]). Cognitive impairment was assessed with the Mini-Mental State Examination (MMSE, [103]) scale. See the test scores **Table 1**.

#### Sample size

Power analysis was informed by a recent study assessing between-group differences in mean volatility estimates in a similar paradigm in healthy individuals with high and low trait anxiety levels ([37]; non-parametric probability of superiority, $\Delta \sim 0.74$, equivalent to a Cohen's d of 0.91, resulting in a minimum of 20 participants per group for 80% power). Differences in frequency-domain amplitude changes during belief updating in that study were associated with a non-parametric effect size of $\Delta \sim 0.735$, equivalent to a Cohen's d of 0.89, resulting in a minimum of 21 participants per group for 80% power. Although larger effects in volatility and neural differences between bipolar patients and their healthy counterparts were anticipated, we recruited 22 BD patients to account for potential variations in effects across computational and neural variables

#### Reward-based motor sequence learning task

Before commencing the main task, participants performed 20 simple sequences of eight finger movements: 1-2-3-4 rightwards and 4-3-2-1 downwards, corresponding to the index, middle, ring, and little fingers on a four-button response box (**Figure 1A**). Each button press produced a specific sound with pitch values E5, F5, G5, A5 (**Figure 1A**). This served as a baseline measure of their fine motor skills.

The primary task had both a familiarisation and test phase. In the familiarisation phase, participants learnt two sequences of four finger presses, each linked to a specific fractal image (**Figure 1A**). The test phase comprised two blocks, totalling 320 trials. In each trial, participants selected between two fractal images, playing the corresponding motor sequence (sequence1, sequence2) for a reward. These sequences, and the corresponding fractals, were implicitly associated with reward values of 5 (rewarded) or 0 points (non-rewarded). The probability of a sequence yielding a reward was reciprocal (p(sequence1|reward) = 1 - p(sequence2|reward)), and shifted pseudorandomly every 26-38 trials. The probability mappings could adopt values of 0.9/0.1, 0.7/0.3, 0.1/0.9, 0.3/0.7, or 0.5/0.5 in each block (**Figure 1B**). Participants received feedback post-trial, indicating 'You earned 5 points' or 'You earned 0 points'. They had a 3000 [± 250] ms window to start and finish the sequence. Feedback was presented 1250 [± 250] ms after sequence completion and remained visible for 1900 [± 100] ms. If participants took over 3300 ms (time out) or made a sequencing error, they were notified of either the delay or mistake, respectively, and garnered no points. Control analyses on the pseudo randomised probabilistic mappings demonstrated that both participant samples experienced the same true volatility. See details in next section.

#### Validation of paradigm and probabilistic structure

The contingencies were generated pseudorandomly in each participant, which could result in both groups experiencing differences in the true volatility by chance. We examined this possibility by comparing the true rate of contingency change in each group: average number of trials where a change takes place across both blocks. A greater number here would denote smaller true volatility( i.e. changes occur less frequently). On average, contingencies changed every 32 trials in both groups (32.3 [SEM 0.25] in HC, 32.9 [ SEM 0.42] in BD), and Bayes factor analysis supported they were equivalent ($BF_{10}$ = 0.61, anecdotal evidence; *P* = 0.1957, non-significant difference).

#### MEG recording and preprocessing

Initial preprocessing of the MEG signals consisted of correcting for head movements, reducing noise, and eliminating bad channels using the temporally extended signal-space separation (tSSS) method (Taulu and Hari, 2009), integrated into the Elekta Neuromag software (Maxfilter, Elekta Neuromag). The settings were adjusted to a sliding window of 10 seconds and a subspace correlation threshold of 0.9.

### Assessing practice effects in general task performance

Bayesian multilevel modelling (BML) estimates approximate posterior probability distributions for model parameters and incorporate accurate uncertainty estimates, even with unbalanced data and small sample sizes [104]. We conducted BML using the R package brms [52, 53], employing informative priors and estimating models by Markov-Chain Monte-Carlo (MCMC) sampling. This process involved drawing 20,000 samples across four chains, discarding the first 1000 iterations of each chain as warm-up. We validated chain convergence using the Gelman–Rubin statistics (R-hat ≤ 1.1; [105]).

For the mIKI analysis and, separately, for the RT analysis, we employed a series of Bayesian multilevel models with a Gaussian distribution. These models included fixed effects, such as Group (BD, HC; with HC as reference) or Trial (1:320), and random effects, capturing the individual variation of subject effects on the intercept and/or slope. Fixed effects refer to effects that are assumed not to vary across subjects, while random effects are considered to vary among individuals (e.g., in their intercept or slope).

Models incorporating Group as a fixed effect facilitated the calculation of the posterior distribution of parameters for the HC group (used as the reference) and the posterior distribution of differences between the BD and HC groups. The most complex model (Model 8 in **Table S1**) included Group, Trial, and their interaction as fixed effects and permitted individual variation in both intercepts and slopes. This approach aimed to reveal how tempo/RT evolved across trials in each participant. For comparative purposes, additional models with varying degrees of complexity were constructed (refer to **Table S1**). Trial-wise outliers in mIKI or RT, defined as values exceeding three standard deviations from the mean, were excluded from the analysis. Moreover, we applied a natural logarithmic transformation to mIKI to normalise its distribution, referred to as log_mIKI (in log-ms). A similar strategy was employed for the analysis of practice effects on RT.

Model comparison was conducted using leave-one-out cross-validation (LOO-CV) with Pareto-smoothed importance sampling [106]. The best-fitting model was identified based on the highest expected log point-wise predictive density (ELPD). We also ensured that the absolute mean difference in ELPD between the two top models (elpd_diff in brms) exceeded twice the standard error of these differences (2se_diff). If elpd_diff was smaller than 2se_diff, we opted for the more parsimonious model.

**Table S1. Bayesian Multilevel Models with a Gaussian distribution assessing practice effects on tempo or RT.** Increasingly complex models were constructed to assess practice effects on tempo (logmIKI, measured in log-ms) or, separately, on RT (logRT, in log-ms). The same models were assessed for each dependent variable, separately, denoted by "*y*" in the table. Models 1, 2, and 3 explain *y* through the fixed effects of group (BD, HC; with HC as the reference group; fixed here denotes an effect assumed not to be changing across subjects) and trial (1:320), or their additive combination, while Model 4 includes their interaction effect. Models 5-7 incorporate the random effect of the intercept by subject, thus modelling how the intercept changes across subjects. The most complex model, Model 8, allows for individual variation in both intercepts and slopes, potentially revealing how individual subjects' tempo/RT changes over trials. Models 4, 7, and 8 enable the assessment of slope differences between BD and HC groups in the practice effect.

| Model # | Model |
|---|---|
| 1 | y ~ 1 + group |
| 2 | y ~ 1 + trial |
| 3 | y ~ 1 + group + trial |
| 4 | y ~ 1 + group*trial |
| 5 | y ~ 1 + trial + (1\|subject) |
| 6 | y ~ 1 + group + trial + (1\|subject) |
| 7 | y ~ 1 + group*trial + (1\|subject) |
| 8 | y ~ 1 + group*trial + (1 + trial\|subject) |
| | The brms family was Gaussian |

In all models, we used a default prior distribution for the intercept, and a normal distribution for each population-level effect (effect of group and trial: normal prior with mean 0 and standard deviation 2; interaction term group x trial: normal prior with mean 0 and standard deviation 1). For the priors on the standard deviation (sd) parameters associated with the group-level effects (individual subjects, trials), we used the default half Student-t prior with 3 degrees of freedom, as recommended by [52]. In Model 8, the prior on the LKJ-Correlation was set to 2.

### Modelling decision-making behaviour using hierarchical Gaussian filters

We employed a validated Bayesian framework with a Hierarchical Gaussian Filter (HGF) to model how input about probabilistic reward outcomes and their change over time is integrated with prior beliefs during learning, resulting in posterior beliefs about the hidden states causing the observed outcomes [23,24]. Drawing inspiration from Behrens et al. [20], in the HGF a sequence of hidden states $\{x_1^{(k)}, x_2^{(k)}, \ldots, x_n^{(k)}\}$—with *k* representing a trial or unit of time—is conceptualised within a generative model consisting of hierarchically coupled Gaussian random walks. These walks are coupled through their variance (inverse precision) and evolve over time.

The HGF generates dynamic and adaptive learning rates, capturing the process of learning under uncertainty in a volatile environment. In this hierarchy, higher levels represent the dynamic structure of the world, and the step size of each random walk is influenced by the state at the level above. Inverting this generative model using variational approximation provides update rules for the temporal trajectories of beliefs held by the agent.

The modelling framework consists of two components: a perceptual model and an observational model. The perceptual model delineates how sensory input—here, reward outcomes—is mapped to the hidden states of the world that generate these inputs. The observational model describes the mapping from the agent's probabilistic representations or beliefs to the produced responses. In other words, the response model accounts for how the decisions of the agent we are observing are derived based on their perception, encompassing both their observations and the inferences drawn from them.

To estimate each participant's individual learning characteristics and belief trajectories during our binary reward-based learning task in a volatile setting, we implemented the three-level perceptual HGF for binary categorical inputs.  At the lowest level, the hidden state $x_1$ corresponds to the binary categorical variable of the experimental stimuli: whether sequence 1 is rewarded in trial $k$ ($x_1^{(k)}= 1$) or not ($x_1^{(k)}= 0$). Beliefs are represented on the second and third levels and modelled as Gaussian distributions. The second level represents the trajectory of participants' beliefs about the contingency between actions (sequence 1 or 2) and their outcomes (rewarded or not), and the third level represents the rate of change in that tendency (volatility). Gaussian belief distributions are represented by their posterior mean ($\mu_2^{(k)}$ , $\mu_3^{(k)}$ for levels 2 and 3 respectively) and posterior variance (uncertainty: $\sigma_2$, $\sigma_3$). Precision is the inverse variance or uncertainty, $\pi_i$ ($i$ = 2 and 3). Belief updating on each level $i$ ($i$ = 2 and 3) and trial $k$ is driven by PEs , modulated by precision ratios, weighting the influence of precision or uncertainty in the current level and the level below:

$$\Delta \mu_i^k = \mu_i^{(k)} - \mu_i^{(k-1)} \propto \frac{\hat{\pi}_{i-1}^{(k)}}{\pi_i^{(k)}} \delta_{i-1}^{(k)} \quad (4)$$

Following equation (4), the expectation of the posterior mean on level $i$, $\mu_i^{(k-1)}$, is updated to its current level $\mu_i^{(k)}$ proportionally to the prediction error of the level below, $\delta_{i-1}^{(k)}$. The influence of PEs is weighted by the ratio of precision values, with the *prediction* (denoted by "^") of the precision of the level below in the numerator and the precision of the current level (inverse uncertainty, $\sigma_i$) in the denominator. Here we assume the prediction from trial $k$-$1$ remains constant until the beginning of the current trial, without drifting to a new value before the agent observes the outcome. Therefore, $\hat{\mu}_i^{(k)} = \mu_i^{(k-1)}$.  See Weber and colleagues [107] for a generalised version of the HGF that explicitly models drifts in the prediction.

In the HGF for binary outcomes, the precision ratio updating beliefs on level 2 in equation (4) is reduced to $\sigma_2^{(k)}$  ($1/\pi_2^{(k)}$), as shown in equation (1) in the main text. Accordingly, the posterior mean of the belief about the action-outcome contingencies is updated via low-level PE about action (stimulus) outcomes, scaled by the degree of informational uncertainty. For level 3, the precision ratio is proportional to the uncertainty about volatility, $\sigma_3^{(k)}$ (inverse precision on level 3: 1/ $\pi_3^{(k)}$).

The HGF perceptual model was coupled with a unit-square sigmoid response model where choice probability is shaped by a free fixed (time-invariant) parameter ζ, interpreted as inverse decision noise: the sigmoid approaches a step function as ζ tends to infinity. The mapping between the predictive probability $m^{(k)}$ for an outcome on trial *k* onto the probability that the individual will choose response 1 or 0, $p(y^{(k)} = 1)$ and $p(y^{(k)} = 0)$ respectively, takes this form (omitting trial index *k* again for simplicity):

$$p(y|m,\zeta)=\left(\frac{m^{\zeta}}{m^{\zeta}+(1-m)^{\zeta}}\right)^{y}\cdot\left(\frac{(1-m)^{\zeta}}{m^{\zeta}+(1-m)^{\zeta}}\right)^{(1-y)} \tag{5}$$

See further detail in Eq. 18 in Mathys et al.[23]. This combination constituted our first perceptual-response model (M1). Next, we used a perceptual 2-level HGF model with volatility fixed to a constant level and coupled it with this unit-square sigmoid response model (M2). Our third model combined the 3-level HGF with a response model where the sigmoid function depends on the trial-wise prediction of log-volatility, $\zeta=e^{-\mu_3^{(k-1)}}$, (M3; [22]). In this observational model, higher estimates of volatility lead to a 'noisier' relationship between beliefs and decision making. As a result, there is an increased likelihood of choosing responses that deviate from predictions. Last, model M4 was constructed similarly to M3 but replaced the free parameter $\omega_2$ with κ[28].

### Selection of priors for HGF models

We selected prior values for our HGF models based on estimates obtained from our data through an ideal observer model. An ideal observer is defined as a model that adopts a range of parameter values aimed at minimising the overall surprise an agent experiences upon receiving a series of inputs (refer to Weber et al. [108] and [71] for different applications of an ideal observer model). This approach was preferred over the use of previously reported prior values for the binary categorical HGF model, primarily because the code in the TAPAS toolbox has undergone modifications and optimisations, leading to improved parameter recovery compared to earlier versions. Consequently, prior values used in previous studies may not be directly applicable to newer model implementations. We used the HGF release v7.1 in Matlab R2020b, and functions 'tapas_ehgf_binary'.

Using ideal model observer models, the group median of the prior values estimated separately for HC and BD were:

$\omega_2$, $\omega_3$: = [-2.5 -0.5] in HC

$\omega_2$, $\omega_3$: = [-2.6 -0.1] in BD

These values were comparable between groups ($BF_{10}$ = 0.4219, 0.7014, support for $H_0$, based on anecdotal and substantial evidence; $P$ = 0.3447, 0.1458, no significant differences). Accordingly, in our main analyses, we selected the median values for [$\omega_2$, $\omega_3$] from the full sample (N = 49), which were [-2.6 -0.3] . For complete details on our prior parameters, see **Table S2**.

In a control analysis employing group-specific priors, we evaluated the consistency of our main computational findings. This analysis, however, should be interpreted with caution due to the comparison of computational variables derived from distinct models for each group. We observed significant group differences in $\omega_2$ ($P_{FDR}$ = 0.0049), as well as in volatility at onset and average volatility ($P_{FDR}$ = 0.0049, 0.0056 respectively). Interestingly, the evidence for differences between groups was stronger than that obtained using common prior settings, with Bayes Factors exceeding 9. This indicates substantial evidence supporting the alternative hypothesis ($H_1$).

**Table S2. Priors (means and variances) on perceptual parameters and starting values of the belief distributions for the winning HGF model M3**

| Prior | Mean | Variance |
|---|---|---|
| $\kappa$ | log(1) | 0 |
| $\omega_2$ | -2.6 | 4 |
| $\omega_3$ | -0.3 | 4 |
| $\mu_2^{(0)}$ | 0 | 0 |
| $\sigma_2^{(0)}$ | log(0.1) | 0 |
| $\mu_3^{(0)}$ | log(1) | 1 |
| $\sigma_3^{(0)}$ | log(1) | 1 |

Quantities are estimated in their native space when they are unbounded, such as $\omega_2$, $\omega_3$. Conversely, quantities with a natural lower bound at zero, like $\kappa$, $\mu_3^{(0)}$, and $\sigma_3^{(0)}$, are estimated in log-space. In the winning HGF model M3, $\omega_2$, $\omega_3$, $\mu_3^{(0)}$, $\sigma_3^{(0)}$ were free parameters ($\kappa$, $\sigma_2^{(0)}$, $\mu_2^{(0)}$ were fixed). The prior variances are in the space in which the corresponding parameter is estimated.

### Assessing motor invigoration

We additionally investigated whether trial-by-trial expectations about the probabilistic action-outcome mapping differentially influenced motor performance in our groups. Considering the association of BD with altered reward sensitivity, even during euthymic phases, we aimed to assess how expectations about reward probability affect motor sequence performance in our task, focusing on performance tempo and RT. Notably, faster sequence completion would expedite the display of the outcome (at 1000 [± 250] ms post-performance).

In the HGF for binary categorical inputs, $\hat{\mu}_2^{(k)}$ represents beliefs about the tendency of the action-outcome contingency. A positive value suggests a higher expectation of reward for sequence 1, and a negative value the reverse (sequence 2 is expected to be more likely to be rewarded). The absolute value, $\left|\hat{\mu}_2^{(k)}\right|$, denotes the strength of predictions about the reward contingency's tendency, regardless of which specific sequence is more likely to be rewarded.

Our hypothesis was that this relationship would exhibit a non-zero (negative) slope, indicating that stronger beliefs about probabilistic contingencies correlate with faster sequence performance. Furthermore, we posited that performance tempo in euthymic bipolar patients might exhibit increased sensitivity to the strength of predictions[4,47], reflected in a comparatively steeper slope than that observed in healthy controls.

To evaluate our motor vigour hypotheses, we constructed a series of BML models with the dependent variable being the logarithm of performance tempo (log_mIKI, in log-ms), in line with our previous work [48]. The most complex model (**Table S3**), incorporated an interaction between group (a categorical variable distinguishing between HC and BD participants) and the centred continuous variable $\left|\hat{\mu}_2^{(k)}\right|$ at trial *k*. This centring step of the continuous predictor is recommended to improve model stability and facilitate the interpretability of the intercept. The interaction term allowed us to determine group-specific associations between predictions and tempo. Additionally, the model included a random effects structure to account for subject-specific variations in intercepts and slopes for the centred predictor, as well as trial-to-trial variations.

Subsequent models of decreasing complexity were constructed by successively omitting random effects or interaction terms. HC served as the reference group for between-group comparisons, providing posterior distributions of differences in slopes and intercepts (see **Table S3** for a full list of models).

For model comparison, we applied the same LOO-CV approach with Pareto-smoothed importance sampling as used in the practice effects analysis. The selection of models was based on the highest ELPD (further details in **General Performance**).

**Table S3. Bayesian Multilevel Models with a Gaussian distribution assessing the effect of strength of predictions on tempo or RT.** Models of decreasing complexity were defined to assess whether a timing variable(tempo [logmIKI] or logRT, represented by *y*) was modulated by the strength of predictions about the tendency of the action-outcome contingency, $\left|\hat{\mu}_2^{(k)}\right|$. This predictor variable was centred and is denoted by *prediction.c* in the table. The most complex model included the random effect of trials, and the random effect of participants on the slope of the timing by prediction association and the intercept.

| Model # | Model |
|---|---|
| 1 | y ~ 1 + prediction.c |
| 2 | y ~ 1 + group*prediction.c |
| 3 | y ~ 1 + prediction.c + (1|subject) |
| 4 | y ~ 1+ group*prediction.c + (1|subject) |
| 5 | y ~ 1+ prediction.c+(1+prediction.c|subject) |
| 6 | y ~ 1+ group*prediction.c + (1+prediction.c|subject) |
| 7 | y ~ 1+ group*prediction.c + (1+prediction.c|subject) + (1|trial) |
| | The brms family was Gaussian |

As for the models listed in **Table S1**, we applied a default prior distribution for the intercept in all models, and a normal distribution for each fixed effect (assuming a constant effect of group and prediction.c: normal prior with mean 0 and standard deviation 2; interaction term group x trial: normal prior with mean 0 and standard deviation 1). For the priors on the sd parameters associated with the random effects (individual subjects, trials), we also employed the default half Student-t prior with 3 degrees of freedom. Models 5-7 include a prior on the LKJ correlation coefficient set to 2.

### Source reconstruction of MEG signals

To source reconstruct MEG signals, integrating planar gradiometers and magnetometers, we used Linearly Constrained Minimum Variance beamforming (LCMV[55]) in the MNE-Python toolbox. We used individual T1-weighted magnetic resonance imaging (MRI) images to establish surface-based cortical divisions for each hemisphere with Freesurfer 6.0 software([56,57]; http://surfer.nmr.mgh.harvard.edu/). MRIs were unavailable for two BD participants; consequently, in those cases, we utilised the fsaverage template brain' files provided by Freesurfer.

We selected the Desikan-Killiany–Tourville atlas (DKT)  atlas, which parcellates the cerebral cortex into 68 distinct anatomical regions[58]. The alignment of MRI and MEG coordinate systems was achieved with an automated procedure within the MNE-Python toolbox (mne.gui.coregistration). This process relied on head position indicator coils and the registered points on the head's surface. Additionally, we ensured that the alignment of three fiducial anatomical points (both preauricular points and the nasion) was accurate across the coordinate systems.

Next, forward modelling was implemented with boundary element conductivity models (BEM) for each participant, with the inner skull surface serving as the chosen volume conductor layout. We then generated a surface-based source space at an "oct6" resolution, which provided 4098 positions (vertices) for every hemisphere with an average adjacent distance of 4.9 mm.

Solving the inverse problem with LCMV beamforming involved computing adaptive spatial filters using a data-covariance matrix for a target interval, selected based on expectations of task effects on neural activity. Informed by prior research indicating modulation of oscillatory activity by pwPE in similar decision-making tasks[37,38], we defined the target interval as 0–1.8 s post-outcome. The noise-covariance matrix for this analysis was estimated from -1 to 0 seconds pre-outcome.

To analyse modulation of activity in the theta (4–7 Hz), alpha (8–12 Hz) and beta (13–30 Hz) frequency bands, the MEG signals were first filtered with a 1–40 Hz band-pass filter, prior to LCMV beamforming. Separately, to assess activity in the gamma frequency range (32–100 Hz), the data were band-pass filtered within 30–124 Hz (as in [37]).

Subsequently, we extracted time courses for individual vertices within cortical labels corresponding to our regions of interest (ROIs), focusing on brain areas consistently associated with decision making, reward-based learning and belief updating under uncertainty. Our selected ROIs included: (1) the ACC, (2) the OFC along with the ventromedial Prefrontal Cortex (vmPFC), (3) the dorsomedial Prefrontal Cortex (dmPFC), (4) the dorsolateral PDC (dlPFC). The ACC and medial PFC are central to emotional regulation, reward processing, and decision making[35,60,61]. Within the medial PFC, the vmPFC is associated with representing reward probability, magnitude, and outcome expectations[62]. Gamma activity in the dmPFC has been correlated with unsigned reward prediction errors during exploration-exploitation behaviour[35]. On the other hand, the dlPFC has been shown to encode belief uncertainty prior to making a decision[63], though some reports link this region to belief updating under uncertainty[64]. Concerning the OFC, this region is crucial for emotional processing and reward/punishment decision making[62], yet different OFC circuits make unique contributions to flexible decision making[109]. Research has associated the medial OFC (mOFC) with encoding reward value, and the lateral OFC (lOFC) with processing nonreward and punishment[110]. In rats, specific OFC circuits have been causally linked to different computations underlying reversal learning[109]. This evidence is particularly pertinent in the context of bipolar disorder as individuals with this condition exhibit impairments in fronto-striatal reward circuitry, including the anterior insula and ventral striatum, but also cortical areas such as the vmPFC, dlPFC[4,10,66,67].

In addition, we included (5) the primary motor cortex (M1) and (6) premotor cortex (PMC) to assess modulation of motor activity during belief updating, and the effect of decision making on motor vigour[68].

Overall, our study's ROIs correspond to **sixteen bilateral labels** in the DKT atlas, encompassing **eight** distinct areas: (1) rostral and caudal ACC (rACC, cACC); (2) lateral and medial OFC (lOFC, mOFC), which include the vmPFC according to some MEG studies[111,112]; but see[113] for a debate on the vmPFC delineation); (3) superior frontal gyrus (SFG), representing the dmPFC and the supplementary motor area (SMA); (4) rostral middle frontal gyrus (rMFG); (5) precentral gyrus (M1), and (6) caudal MFG (cMFC).

The representative time series for each label were derived using the 'PCA flip' method in MNE-Python. This method employs singular value decomposition (SVD) on the time courses associated with each vertex within a specific brain region or label. Its primary function is to extract the first right singular vector, representing the direction of maximal power in each source. Following this extraction, the singular vector undergoes scaling and sign-flipping, which then yields the time course for the region of interest (ROI). Although the 'flip' operator was not relevant for our time-frequency (TF) analysis, it was essential for preparing the source-reconstructed time series for subsequent connectivity analysis.

### Convolution modelling of time-frequency responses during outcome processing

We employed a general linear model (GLM) to investigate the frequency-domain, trial-by-trial amplitude changes associated with belief updating and processing uncertainty following the presentation of the outcome. In the winning HGF model, the trajectories of beliefs at levels 2 and 3 are updated proportionally to the magnitude of pwPEs: low-level pwPEs about the action-outcome contingencies, termed $\varepsilon_2$, and high-level pwPEs about the environmental volatility, $\varepsilon_3$. As the sign of pwPE changes updating level 2 is arbitrary in the binary categorical HGF, previous studies have opted for using the unsigned $|\varepsilon_2|$ as a regressor in GLM analyses of neural activity[37,38]. Variable $|\varepsilon_2|$ represents precision-weighted Bayesian surprise about reward outcomes, independent of whether they are related to sequence 1 or 2. Given the collinearity of $\varepsilon_3$ and $|\varepsilon_2|$(in each participant, the trajectories were highly correlated, Pearson $R$ in range 0.52–0.94, $P < 10^{-10}$ in all cases), we selected $|\varepsilon_2|$ as our primary parametric regressor, modulating induced oscillatory activity following the outcome presentation. Additional parametric regressors included informational uncertainty, $\sigma_2$, and level 3 uncertainty, $\sigma_3$, as the relevant (inverse) precision terms for the GLM. Furthermore, our GLM incorporated discrete regressors coding for win and lose outcomes, and error trials (performance errors or timeouts).

We implemented the GLM on the time series of induced responses over multiple frequencies using a linear convolution model developed by Litvak and colleagues[69]. This approach, an adaptation of the classical GLM used in fMRI analysis for time-frequency data, has been successfully employed in previous work to identify frequency-resolved neural correlates of belief updating, precision, and predictions during decision-making and perceptual learning[36,37,38]. It facilitates assessing the modulation of TF responses on a trial-by-trial basis by a specific explanatory regressor while controlling for the effects of other regressors included in the model.

This outcome-locked convolution model was solved in the source space after applying LCMV to the time series of concatenated epochs of MEG data (see previous section). We tested the hypothesis that bipolar patients exhibit changes in gamma and concomitant alpha/beta activity during the encoding of pwPE signals. Additionally, we hypothesised that euthymic BD individuals would show alterations in alpha/beta oscillatory activity during the representation of precision weights.

To conduct the convolution GLM, we estimated standard TF representations of the source-level time series using Morlet wavelets. TF spectral power was extracted between 4 and 100 Hz and transformed to amplitude, following ref. [69]. For the theta (4–6 Hz), alpha (8–12 Hz), and beta (13–30 Hz) frequency ranges, we used 5-cycle wavelets shifted at every sampled point in bins of 2 Hz. For gamma-band activity (32–100 Hz), 7-cycle wavelets sampled in steps of 2 Hz were employed. To represent any physiological response shape, the convolution GLM approach considers a generic basis set (e.g., sines and cosines). Regressors were constructed by convolving a set of basis functions with a set of input functions representing the events of interest. We chose a 20th-order Fourier basis set (comprising 40 basis functions: 20 sines and 20 cosines). Each discrete and parametric regressor was convolved with this set. This setting allowed the GLM to resolve modulations of TF responses up to approximately 8.7 Hz (20 cycles / 2.3 seconds, or ~115 ms). See further details in the original methodological publication[69].

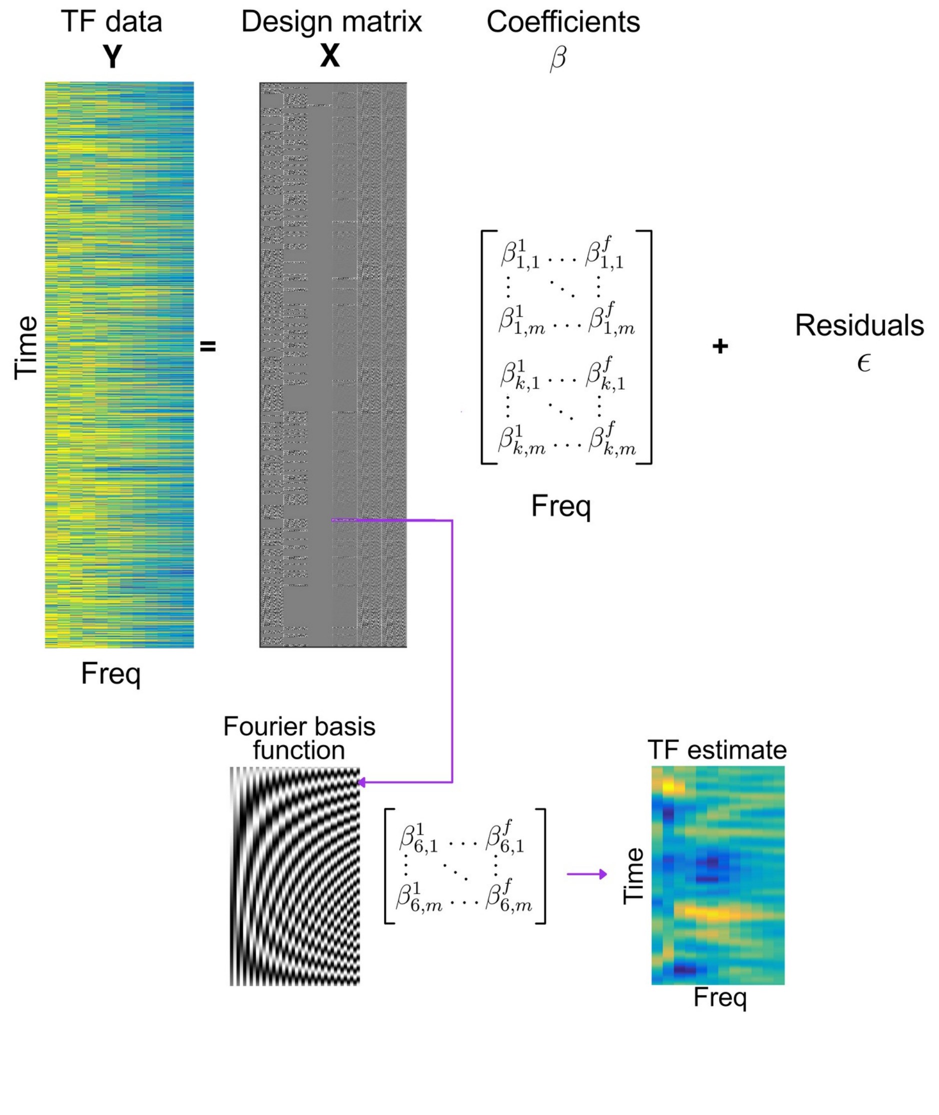

**Figure S1. Convolution General Linear Model (GLM).** This figure illustrates the standard pseudo-continuous time-frequency (TF) representations of the source-reconstructed MEG signal (**Y**), estimated using Morlet wavelets. In the GLM framework, the signal **Y** is modelled as a linear combination of explanatory variables or regressors (contained in matrix **X**), modulated by regression coefficients (**β**), with an additional noise term (**ϵ**). The design matrix **X**, depicted in the lower left inset, is constructed by convolving Fourier functions (m = 20; comprising 20 sine and 20 cosine functions) with input functions that represent the timings and values of relevant discrete or parametric events. For this example, six regressors were used (from left to right in the figure): Outcome Win, Outcome Lose, Outcome No Response, and absolute precision-weighted prediction error (pwPE) on level 2, $\sigma_2$, and $\sigma_3$, each defined over time. The solution to the convolution GLM yields response images (represented by the TF estimate in arbitrary units in the figure) as a combination of the basis functions *m* and the regression coefficients $\beta_i$ for each specific regressor type *i*, across frequencies *f* and basis functions *m*. Consequently, the convolution GLM effectively estimates deconvolved TF responses (shown as the TF estimate in the rightmost bottom image) to event types and their associated parametric regressors. See details in Litvak and colleagues[69].

### Frequency-resolved connectivity

To assess functional connectivity between frequency-resolved activity in our reconstructed sources, we employed a robust metric of directed information flow between time series: time-reversed Granger causality (TRGC,[73,74]. This metric builds upon Granger causality (GC), a statistical method used to determine directional information flow between signals[114]; frequency domain: [115]). GC operates on the principle that, if information from the past of signal *x(t)* improves the prediction of the present of signal *y(t)* relative to what can be predicted by the past of *y(t)* alone, then *x(t)* is said to 'Granger-cause' *y(t)*.

Simulation work demonstrated that GC is influenced by volume conduction and source leakage, affecting its robustness[73]. To address these limitations, TRGC was developed. Haufe and colleagues[73] proposed using time-reversed data as surrogates for statistical testing when assessing GC between pairs of time series in a multivariate setting. Spurious directional flow between time series, explainable by volume conduction and corresponding to zero-lag directional influences, is expected to persist in the time-reversed analysis. By correcting the GC coefficients with those from the time-reversed GC analysis, an unbiased estimate of directional flow is obtained, one not influenced by volume conduction or source leakage. This approach has been validated through simulation work, establishing TRGC as a robust metric for inferring the correct directionality of information flow. In addition to mitigating the influence of hidden common drivers, TRGC can mitigate the influence of measurement noise[116,117].

In our study, we estimated frequency-domain TRGC to assess directional information flow between our 16 ROIs (8 bilateral ROIs). Following the recommendations of Pellegrini et al. [74], we used the source-reconstructed time series in our ROIs obtained through LCMV beamforming. This process included a dimensionality reduction step with PCA/SVD to select a singular vector associated with the direction of maximal power in each source (refer to the *Source reconstruction of MEG signals* section). We focused the TRGC analysis on the alpha and beta frequency ranges, and assessed the directionality of functional coupling during the 0.5–1 s interval of outcome processing for trials with larger unsigned pwPEs updating beliefs at level 2 (median split per participant). The TRGC analysis was implemented using the ROIconnect plugin for EEGLAB in MATLAB from ref. [74], available at https://github.com/sccn/roiconnect. We adapted the code for application to our LCMV output obtained in MNE-python.

#### Statistical analyses

To address the multiple comparisons problem, which arises in contexts such as several post-hoc analyses, we control the false discovery rate (FDR) using an adaptive linear step-up procedure [118], set to a level of q = 0.05. This provides an adapted threshold p-value ($P_{FDR}$). In the case of pairwise statistical analyses, we provide estimates of non-parametric effect sizes for pairwise comparisons, along with associated bootstrapped confidence intervals [75,76]. The between-group effect sizes are estimated as the probability of superiority (Δ).

### Supplemental Results

#### Demographics

Age and sex distribution were comparable across groups (Age: $P$ = 0.4921, no-significant differences; $BF_{10}$ = 0.3472, providing anecdotal evidence for $H_0$; Sex: Chi-squared statistic, $\chi^2$= 1.6559, *df = 1*, $P$ = 0.1849, no significant; $BF_{10}$ = 1.102, no evidence for $H_0$ or $H_1$).

BD and HC groups did not significantly differ in their anxiety or depression scores (BDI score, $P_{FDR}$ = 0.1038, $BF_{10}$ = 0.8668, anecdotal evidence for $H_0$; State-Trait Anxiety Inventory, state subscale: $P_{FDR}$ = 0.4571, $BF_{10}$ = 0.3833, anecdotal evidence for $H_0$; HADS, depression subscale: $P_{FDR}$ = 0.4003, $BF_{10}$ = 0.3824, anecdotal evidence against group differences; anxiety, $P_{FDR}$ = 0.64, $BF_{10}$ = 0.31, substantial evidence for the null hypothesis). Altman's Mania scores were also simlar between groups ($P_{FDR}$ = 0.2040, $BF_{10}$ = 0.5652). Refer to **Table 1.**

Regarding cognitive abilities, we identified significant group differences in the second part of the TMT, assessing executive functioning (independent-sample permutation test, $P_{FDR}$ = 0.0022; non-parametric effect size estimator: Δ = 0.80, CI = [0.63, 0.92]). By contrast, BD and HC participants performed similarly in the less challenging first TMT part ($P_{FDR}$ = 0.2216, $BF_{10}$ = 0.5296, providing anecdotal evidence for $H_0$). Differences in Wisconsin Card Sorting Test were not significant after FDR control ($P$ = 0.0364; non-significant, as the adjusted significance level was 0.0022; $BF_{10}$ = 1.6560, anecdotal evidence for $H_1$). Last, both samples had similar MMSE scores ($P$ = 0.8678, $BF_{10}$ = 0.2875, providing substantial evidence for the null hypothesis).

### Altered reward-based decision dynamics in bipolar disorder during euthymia

BD showed changes in the expression of win-stay/lose-shift behaviour relative to HC(see main text). Notably, there was no between-group difference in their decision to switch strategies post-loss, based on anecdotal evidence (lose-shift rate: $P$ = 0.0966, non-significant difference; $BF_{10}$ = 0.8905; **Figure 2d**). Overall, when considering all types of switches, BD individuals changed their responses more than healthy participants (total switch rate, 0.16 [0.019] in HC, 0.23 [0.049] in BD; $P_{FDR}$ = 0.0160; $\Delta$ = 0.70, CI = [0.55, 0.85]). Both groups committed errors at a similar rate during the task performance (around 1%, $P$ = 0.1482, non-significant difference; $BF_{10}$ = 0.6867, anecdotal evidence for $H_0$).

We next examined these behavioural results further using the HGF as a computational framework. Applying Bayesian model selection to models M1-M4 in the total sample, we found that the M3 model outperformed the other models across all subjects (exceedance probability, *Pexc* = 0.77; expected frequency, *Freqexp* = 0.45; **Table S4**). Model M3 was also the winning model separately for HC group (*Pexc* = .52, *Freqexp* = 0.41), and BD group (*Pexc* = .70, *Freqexp* = 0.43). In the M3 response model, decisions depend on dynamic trial-by-trial estimates of log-volatility. A heightened expectation of environmental volatility on the current trial results in higher decision noise, implying a noisier belief-response mapping. Conversely, when the environment is anticipated to be more stable, the coupling between the current belief and the ensuing decision becomes more deterministic. This model also allows for individual differences in $\omega_2$, the tonic volatility of action-outcome associations, as well as time-invariant volatility on the third level, $\omega_3$; free model parameters also include the initial values $\mu_3^{(0)}$ and $\sigma_3^{(0)}$.

**Table S4. Bayesian model selection (BMS).** Log model evidence was used for model comparison using random-effects Bayesian model selection[119]. The columns display the exceedance probability, *Pexc*, and expected frequency, *Freqexp*, for models M1-M4. Model M3 best explained the data in our total sample (N = 49), and in each group separately (HC, N = 27; BD, N = 22). BMS was implemented using code from the MACS toolbox[120].

| Model, Sample | Pexc | Freqexp |
|---|---|---|
| M1, HC+BD | 0 | 0.0239 |
| M2, HC+BD | 0.1920 | 0.3226 |
| M3, HC+BD | **0.7701** | **0.4484** |
| M4, HC+BD | 0.0379 | 0.2050 |

| | | |
|---|---|---|
| M1, HC | 0 | 0.04 |
| M2, HC | 0.44 | 0.38 |
| M3, HC | **0.52** | **0.41** |
| M4, HC | 0.04 | 0.17 |
| M1, BD | 0 | 0.05 |
| M2, BD | 0.12 | 0.26 |
| M3, BD | **0.7** | **0.43** |
| M4, BD | 0.19 | 0.26 |

Simulations conducted to assess the accuracy of parameter estimation in the winning HGF model M3 demonstrated that parameters $\omega_2$ and $\mu_3^{(0)}$ were estimated with high accuracy. By contrast, $\omega_3$ and $\sigma_3^{(0)}$ were recovered with less accuracy, as previously reported[27,70]. See **Figure S2** for details. Specifically, we simulated behavioural responses of 50 agents across six different values of $\omega_2$ and, separately, six values of $\omega_3$, using the input observed for one of our participants (ID 47). This resulted in 300 simulated agents for each individual $\omega_2$ or $\omega_3$ value. To determine the estimation accuracy of parameters $\mu_3^{(0)}$ and $\sigma_3^{(0)}$, we conducted similar simulations with 50 agents across six values each for $\mu_3^{(0)}$ and $\sigma_3^{(0)}$.

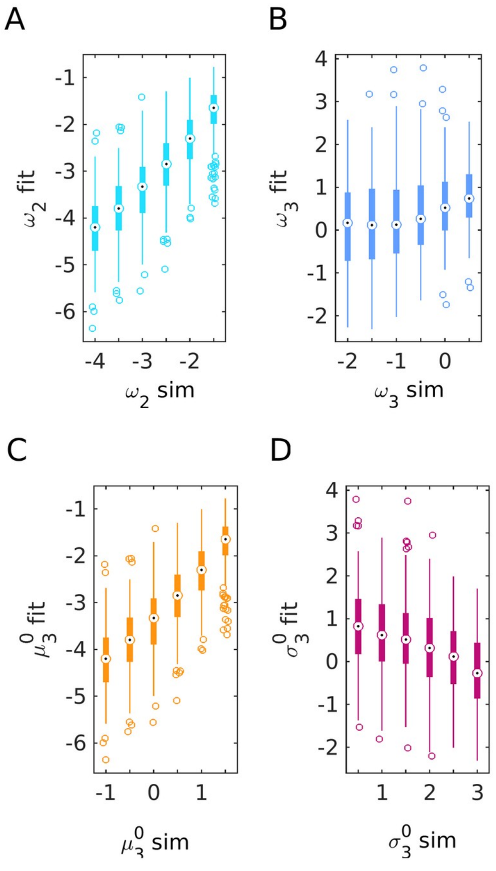

**Figure S2. HGF parameter recovery.** This figure illustrates the estimation of HGF parameters using the input observed from one of our participants (ID 47). Panels **a-d** display boxplots (showing the median, 25th, and 75th percentiles) of parameter estimation results for $\omega_2$ (a), $\omega_3$ (b), $\mu_3^{(0)}$(c), and $\sigma_3^{(0)}$(d). The x-axis represents the parameters set in the simulated responses (labelled "sim"), and the y-axis shows the corresponding estimated values of those parameters (labelled "fit"). Parameters $\omega_2$ and $\mu_3^{(0)}$ were estimated with high accuracy, indicated by a significant correlation between simulated and estimated (fit) values: Pearson $R = 0.8158$, $P \ll 1 \times 10^{-6}$ for $\omega_2$, and $R = 0.8049$, $P \ll 1 \times 10^{-6}$ for $\mu_3^{(0)}$. By contrast, parameter $\omega_3$ showed less accuracy in estimation: $R = 0.2761$, $P \ll 1 \times 10^{-6}$, as did $\sigma_3^{(0)}$: $R = -0.0093$, $P = 0.6920$. The prior values for $\omega_2$, $\omega_3$ , $\sigma_3^{(0)}$, and $\mu_3^{(0}$ used in the configuration file for estimating each parameter from the simulated responses, as defined in **Table S2** for the best fitting model M3, were: $\omega_2 = -2.6$, $\omega_3 = -0.3$ (with a variance of 4 for both); $\mu_3^{(0)} = 1$, $\sigma_3^{(0)} = 1$ (with a variance of 1 for both in log space).

Using the winning model to assess between-group differences in the model parameters $\omega_2$, $\omega_3$, $\mu_3^{(0)}$, as well as in the mean log-volatility estimate over trials, $\mu_3$, we reported in the main text significant differences in $\omega_2$, $\omega_3$, $\mu_3^{(0)}$, $\mu_3$. In particular, initial volatility and its trial-average were higher in BD than in HC, while $\omega_2$ was lower. Parameter $\omega_2$ influences the coupling of beliefs between levels 2 and 3 through changes in environmental uncertainty, defined as $\exp(\kappa\mu_3^{(k-1)} + \omega_2)$[24]. Thus, despite BD patients perceiving the environment as more volatile, their belief updates about reward contingencies were not as influenced by this expectation compared to the HC group. As expected, log-volatility estimates correlated with the total switch rate, accounting for response stochasticity (**Figure S3**). Last, there were no significant differences in tonic volatility on level 3, $\omega_3$ ($P = 0.0730$, anecdotal evidence suggesting group differences, $BF_{10} = 1.09$).

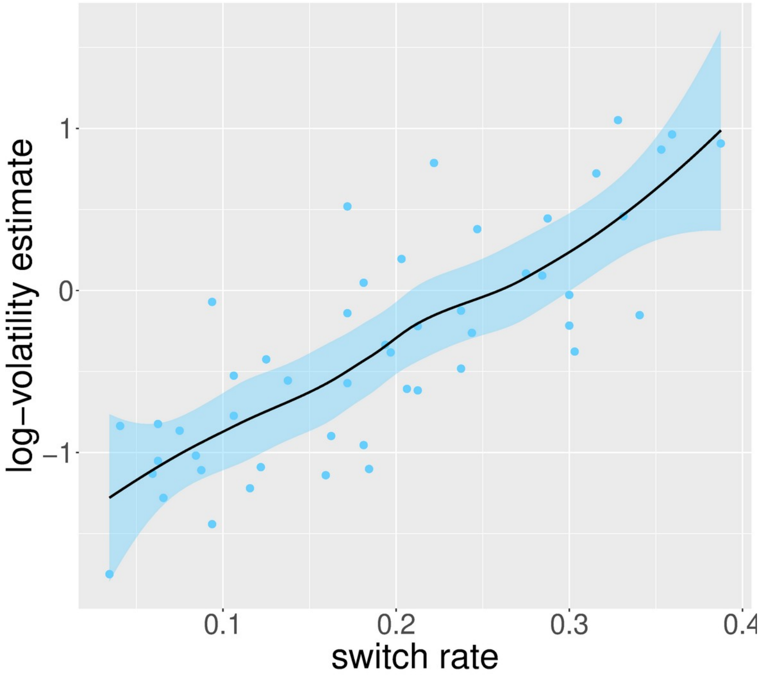

**Figure S3. Non-parametric rank correlation between phasic log-volatility and total switch rate.** There was a significant positive Spearman rank correlation between the total switch rate and the average phasic log-volatility, $\mu_3$ (N = 49; ρ = 0.79 [95% CI: 0.65, 0.88], $P = 1.7 \times 10^{-11}$). Individual participant data points are depicted as blue circles. The black line indicates the non-linear fit estimated using the Locally Weighted Scatterplot Smoothing (LOESS) method via the geom_smooth() function in R, employing the ggplot2 library. The graphic also features the 95% confidence interval as a shaded blue area.

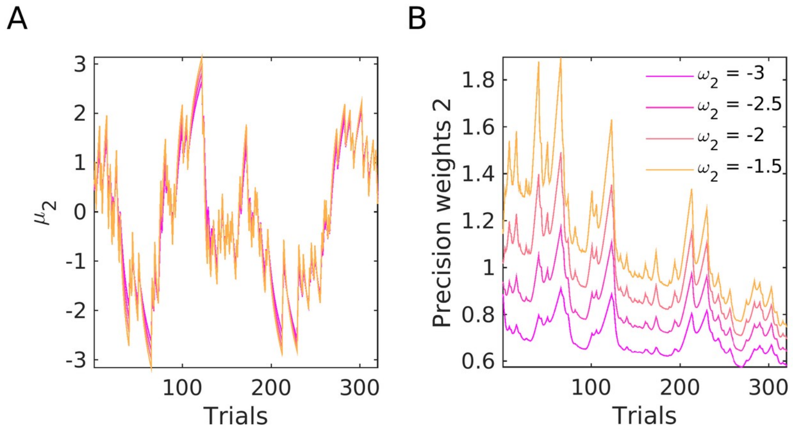

**Figure S4. Simulated belief trajectories and precision weight terms on level 2. a.** In the winning model, M3, belief trajectories for $\mu_2$ were simulated using input data from one participant (ID 47), with priors set at $\mu_2^{(0)} = 0$, $\sigma_2^{(0)} = 0.1$, $\mu_3^{(0)} = 1$, $\sigma_3^{(0)} = 1$, $\kappa = 1$, $\omega_3 = -0.5$, but with $\omega_2$ being modulated. This parameter, $\omega_2$ , represents the tonic part of the variance in the Gaussian random walk for $x_2$ and modulates the learning rate about response outcomes at the lowest level. We demonstrate that decreasing $\omega_2$ (represented by changes from orange to magenta lines) is associated with smaller update steps in the belief about the action-outcome contingency tendency, $\mu_2$. **b.** In line with (a), decreasing $\omega_2$ also reduces the estimation uncertainty about the reward tendency, $\sigma_2$. Since $\sigma_2$ represents the precision weights term scaling belief updating at level 2, this simulation suggests that lower tonic volatility, $\omega_2$, has a slowing effect on belief updating about the action-outcome contingency tendency .

### Correlations between HGF parameters and residual symptoms

To assess associations between residual symptoms and HGF parameters, we conducted non-parametric Spearman rank correlations, given that residual symptoms are represented as ordinal data. Utilising the R library “correlation” and the method “spearman”, we identified a significant positive correlation between volatility estimates and trait anxiety (**Figure S5A**), as well as a negative association between mania and estimation uncertainty $\sigma_2$ (**Figure S5B**). However, no significant correlation was found between depression and $\sigma_2$, as detailed in the main document.

Exploring whether the significant correlations in the BD sample extended to the full sample (N = 49 participants), we observed that log-volatility and trait anxiety were indeed significantly correlated ($\rho$ = 0.31, 95% confidence interval [0.02, 0.55], $P_{FDR}$ = 0.033). Mania scores and $\sigma_2$ were, however, not associated in the full sample (N = 48; $\rho$ = -0.17 [-0.42, 0.08] , $P$ = 0.740, $BF_{10}$ = 0.740).

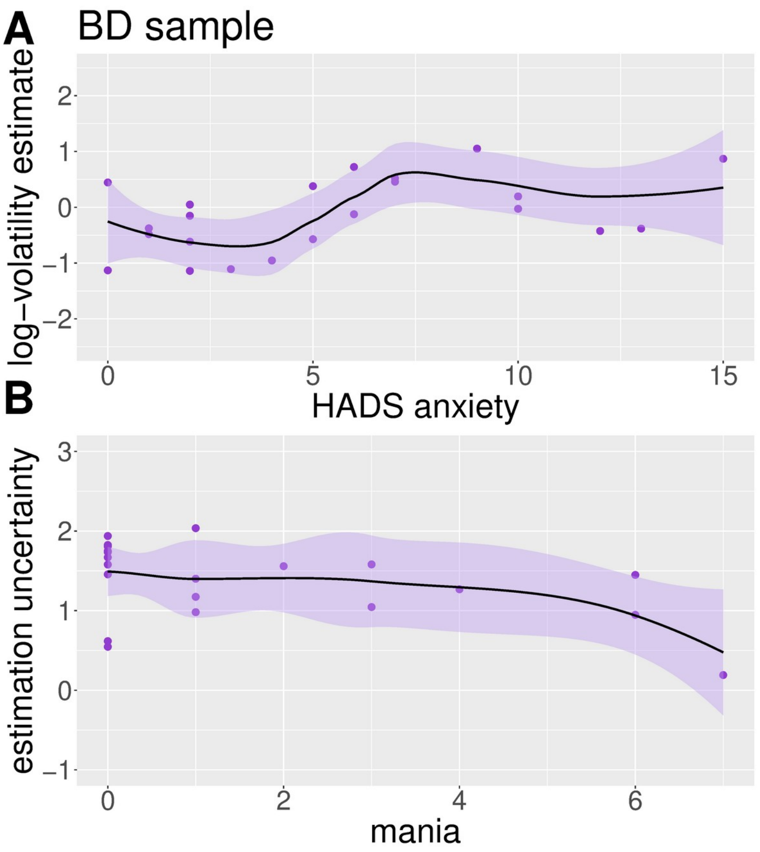

**Figure S5. Non-parametric rank correlation between residual symptoms and relevant HGF variables. a.** The Spearman rank correlation between trait anxiety levels and the mean expectation on log-volatility was positive and significant (N = 22; $\rho$ = 0.46, 95% confidence interval [0.04, 0.75], $P_{FDR}$ = 0.030). The purple circles denote individual BD values, while the black line illustrates the non-linear fit estimation (LOESS). The 95% confidence interval is represented as a shaded purple area. **b.** Same as a but variables $\sigma_2$, estimation uncertainty, and mania scores in BD. There was a significant negative association between these variables (N = 21; $\rho$ = -0.46 [-0.75, -0.02], $P_{FDR}$ = 0.037; mania scores were not available in one patient).

### Behavioural results relative to an ideal Bayesian observer

To assess whether deviations in decision-making behaviour in BD could be ascribed to larger departures from the expected learning patterns of an ideal Bayesian observer than in HC, we simulated the behavioural responses of agents observing the input of each participant in both cohorts. Using the same priors as in the primary modelling analysis, we extracted the time series of behavioural responses in each agent and computed the simulated win rates, win-stay and lose-shift rates (**Figure 2cd**).

First, we confirmed that these rates were comparable between two groups of ideal observers, representing HC and BD. This demonstrated that the pseudorandomised task structure was balanced across groups ($P$ = 0.2496, 0.5663, 0.07; $BF_{10}$ = 0.5200, 0.3295, 0.5450: substantial and anecdotal evidence supporting no differences for win rates, win-stay and lose-shift rates, respectively).

Subsequently, we found that both win rates and win-stay rates were significantly lower in the participant groups compared to the simulated values, as expected. This was the case for both HC and BD (HC: $P_{FDR}$ = 0.0004 for win rate, 0.0002 for win-stay rate; BD: $P_{FDR}$ = 0.0002 for both metrics). Lower win-stay rates in our participant groups than in the simulated agents indicated that they were more likely to change after securing a win on a trial.

In terms of lose-shift behaviour, healthy participants exhibited rates akin to those of ideal Bayesian observers (*P* = 0.1076, with the difference being non-significant; $BF_{10}$ = 0.6775, offering anecdotal evidence for $H_0$). Conversely, BD patients demonstrated a more pronounced tendency to shift after a loss compared to their ideal Bayesian learner counterparts, which was significant ($P_{FDR}$ = 0.0004).

### Similar timing of actions during motor decision-making in euthymic bipolar and healthy participants

Baseline motor performance in bipolar participants was slower compared to healthy individuals (performance tempo assessed as average inter-key press-interval, IKI, in ms: 266 [12.2] ms in HC, 323 [19.1] ms in BD, *P* = 0.02; $\Delta$ = 0.73, CI = [0.54, 0.87]). By contrast, during the primary motor decision-making task, both groups performed motor sequences at similar tempi, approximately three key presses per second (mean IKI, mIKI: 339 [16.6] ms in HC, 354 [18.2] ms in BD, *P* = 0.6223; $BF_{10}$ = 0.33, indicating substantial evidence against group differences). Regarding reaction time—the interval before participants initiated the sequence—both groups were comparable, though this is based on anecdotal evidence (mean RT 501 [20.1] ms in HC, 553 [32.2] ms in BD, *P* = 0.2262; $BF_{10}$ = 0.55).

We subsequently examined practice effects to assess improvements in performance tempo and RT across trials. For both of our timing dependent variables (DV), log-mIKI and log-RT (**Figure 3a, d**), LOO-CV identified the most complex model, model 8 (**Table S1**) as the best fit (**Table S5**). For log-mIKI, the absolute mean difference in ELPD between the model 8 and the second best fitting model (model 7) was 763.8061 and the standard error of the differences (se_diff) was 46.66605 (elpd_diff > 2*se_diff). When ELPD differences between two models are larger than four, and if the number of observations is > 100, and the model is moderately well specified, then the standard error is a good estimate of the uncertainty in the difference between models[116,121].

This model described changes in motor performance (either log-mIKI or log-RT) across trials for the reference group of healthy controls, captured by the fixed effect of the trial. The modulation of these practice effects by group was represented by the interaction term group*trial. This interaction reflects the differences in slopes between BD and HC groups, illustrating how changes in timing across trials varied between groups. In addition, the inclusion of the random effects term (1 + trial|subject) in the structure of this model allows for both the intercept (baseline DV) and the slope (change in DV across trials) to vary by subject, directly capturing the individual differences in how practice affects tempo or RT.

Posterior predictive checks indicated that the best model for tempo demonstrated robust predictive accuracy across the range of the DV **(Figure S6a)**. The predictive strength of the best model for RT was, however, lightly diminished (**Figure S6b). Table S5** presents a summary of the posterior point estimates for the winning model.

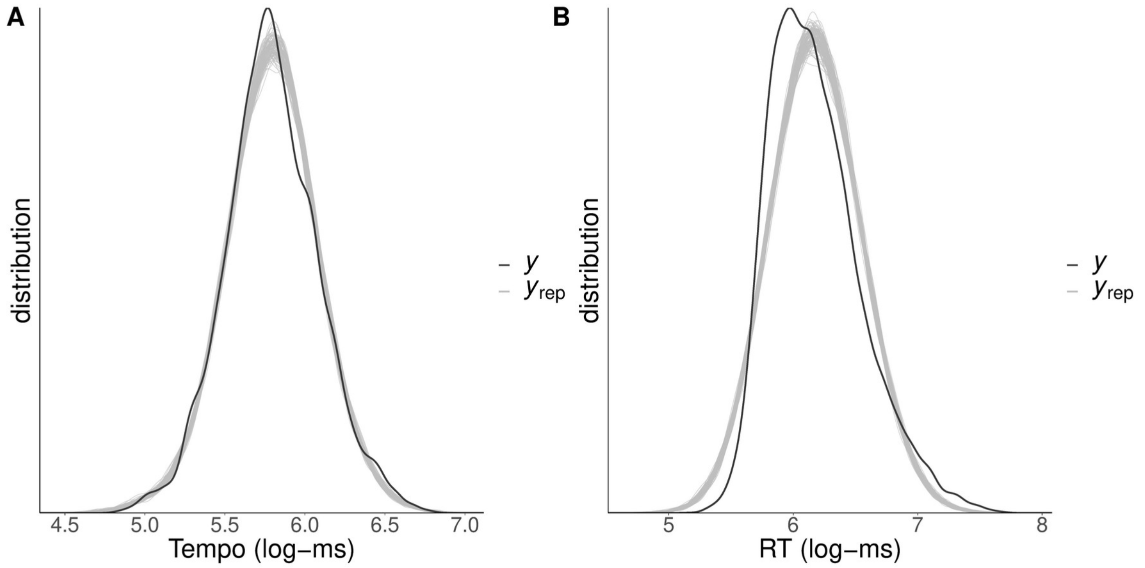

**Figure S6. a.** Illustration of the posterior predictive checks where the distribution of the observed outcome variable (y, in our case performance tempo, log-transformed) is compared to simulated datasets ($y_{rep}$) from the posterior predictive distribution (100 draws). **b**. same as a, but for log-RT.

**Table S5. Summary parameter estimates for the winning Bayesian multilevel model assessing practice effects.**

| Dependent Variable | Parameter (Population-level effects) | Estimate | l-95% CI | u-95% CI | R-hat |
|---|---|---|---|---|---|
| Performance tempo (log-mIKI, in log-ms) (model8) | | | | | |
| | Intercept | **5.79014** | **5.68824** | **5.89292** | 1.02 |
| | Group (BD - HC) | 0.08536 | -0.04126 | 0.21718 | 1.02 |
| | Trial | **-0.00008** | **-0.00012** | **-0.00003** | 1.00 |
| | Group:Trial | **-0.00022** | **-0.00029** | **-0.00016** | 1.00 |
| Reaction time (log-RT, in log-ms) (model8) | | | | | |
| | Intercept | **6.21087** | **6.11646** | **6.30298** | 1.00 |
| | Group (BD - HC) | 0.11165 | -0.02092 | 0.24376 | 1.00 |

| | | | | |
|---|---|---|---|---|
| Trial | **-0.00037** | **-0.00063** | **-0.00013** | 1.00 |
| Group:Trial | -0.00031 | -0.00067 | 0.00005 | 1.00 |

Estimate = posterior mean; CI = credible interval based on quantiles. Gelman-Rubin statistics demonstrate chain convergence (R-hat < 1.1; [105]).

Bayesian multilevel modelling indicated that both groups started with comparable timing (log-mIKI and log-RT: the 95% credible interval, CI, for the difference in intercept estimates overlapped with zero). In addition, for both DV there was a credible fixed effect of trial number on changes in timing performance (HC as reference group; negative slope: the 95% CI of the estimated slope for HC did not include zero). This means that performance timing (mIKI and RT) improved across trials in HC. Notably, the BD group exhibited more rapid improvements in their performance tempo across trials (steeper slope: **Figure 3bc**) than their healthy counterparts (the 95% CI for the estimate of slope differences did not encompass zero). This was not the case for RT (no credible effect for slope differences; **Table S5**).

In conclusion, while baseline motor performance varied between groups, the timing of actions during the main motor decision-making task was similar across both groups. Nevertheless, BD participants displayed a more pronounced slope for practice effects on tempo.

### Expectation about reward probability invigorates motor performance similarly in euthymic bipolar and healthy participants

We next asked whether decision-making differences between bipolar and healthy participants lead to distinct motor invigoration effects. In a similar task, we previously showed that the strength of predictions about action-outcome contingencies is associated with faster performance tempo in sequences of finger movements[48]. BD individuals, including those in euthymic phases, have been reported to mobilise more effort, confidence, and energy while expecting and receiving rewards[47]. Beyond the observed practice effects in BD patients compared to HCs, we therefore considered that BD individuals might exhibit altered invigoration effects on a trial-by-trial basis, influenced by their expectations about the action-outcome mapping.

BML modelling assessing the association between log-mIKI and $\left|\hat{\mu}_2^{(k)}\right|$ revealed that three of the tested models equally well explained the data, as indicated by LOO-CV (models 5–7: **Table S3**). Since the difference in ELPD was smaller than twice the standard error (2*se_diff) for these models, we selected the most parsimonious model, model 5. The ELPD difference to the next best model met our criterion elpd_diff > 2*se_diff ( 29.641673 > 2 * 9.086587). Model 5, which did not include group differences or trial-specific random effects, effectively explained performance tempo through the strength of participants' predictions about reward contingencies ($\left|\hat{\mu}_2^{(k)}\right|$), while also accounting for individual variability on the slope and intercept (subject included to model random effects). Posterior point estimates for the effects in the best model are shown in **Table S6**.

The posterior predictive checks demonstrated that the observed outcome variable y (log-mIKI) overlapped well with the simulated datasets $y^{rep}$ from the posterior predictive distribution (**Figure S7a**).

**Table S6. Summary parameter estimates for the winning Bayesian multilevel model assessing the effect of strength of predictions on timing performance.**

| Dependent Variable | Parameter (Population-level effects) | Estimate | l-95% CI | u-95% CI | R-hat |
|---|---|---|---|---|---|
| Performance tempo (log-mIKI, in log-ms) (model5) | | | | | |
| | Intercept | **5.78357** | **5.71471** | **5.85564** | 1.00 |
| | Predictions | **-0.00753** | **-0.01452** | **-0.00046** | 1.00 |
| Reaction time (log-RT, in log-ms) (model5) | | | | | |
| | Intercept | **6.14696** | **6.06355** | **6.23095** | 1.00 |
| | Group (BD-HC) | 0.06542 | -0.05844 | 0.18618 | 1.00 |
| | Predictions | 0.01003 | -0.00556 | 0.02522 | 1.00 |
| | Group:Predictions | -0.00799 | -0.03083 | 0.01492 | 1.00 |

Estimate = posterior mean; CI = credible interval based on quantiles. Gelman-Rubin statistics demonstrate chain convergence (R-hat < 1.1). The predictor "Predictions" denotes the centred values of the strength of participants' predictions about reward contingencies ($\left|\hat{\mu}_2^{(k)}\right|$).

Model 7, the most complex for RT, emerged as the best based on ELPD criteria. Although this model showed relatively good posterior predictive accuracy, it was slightly lower than the model for tempo. It revealed no credible effects of the strength of predictions on log-RT among participants, nor were there group effects, as the 95% CI for these effects included zero (**Table S6; Figure S7b**)

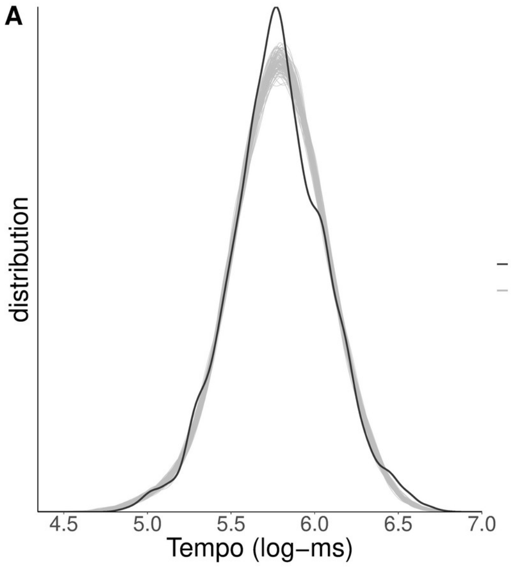

**Figure S7. a.** Illustration of the posterior predictive checks for log-mIKI BML model 5, where the distribution of the observed outcome variable (y, in our case performance tempo, log-transformed) is compared to simulated datasets ($y_{rep}$) from the posterior predictive distribution (100 draws). **b**. same as a, but for log-RT.

### Neural representation of pwPE updating beliefs about the action-outcome contingencies

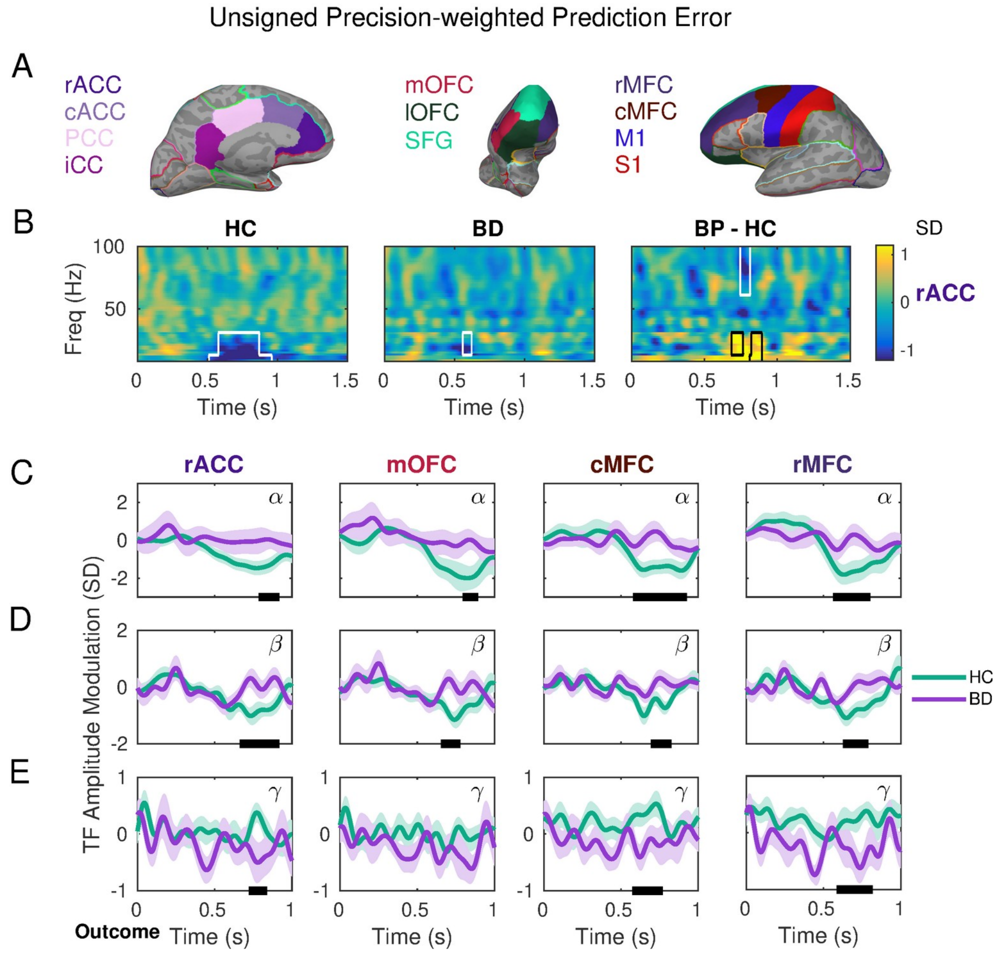

**Figure S8. Attenuated gamma increase and alpha-beta suppression during encoding unsigned precision-weighted prediction errors about stimulus outcomes in bipolar disorder.** Same as **Figure 4** but in additional ROIs where between-subject differences were observed after FWER control at 0.025: rACC, mOFC, caudal and rostral MFG. Labels denote the rostral anterior cingulate cortex, rACC; caudal ACC, cACC; superior frontal gyrus, SFG; lateral and medial orbitofrontal cortex, lOFC and mOFC; primary motor cortex, M1; caudal and rostral middle frontal gyrus, cMFG, rMFC.

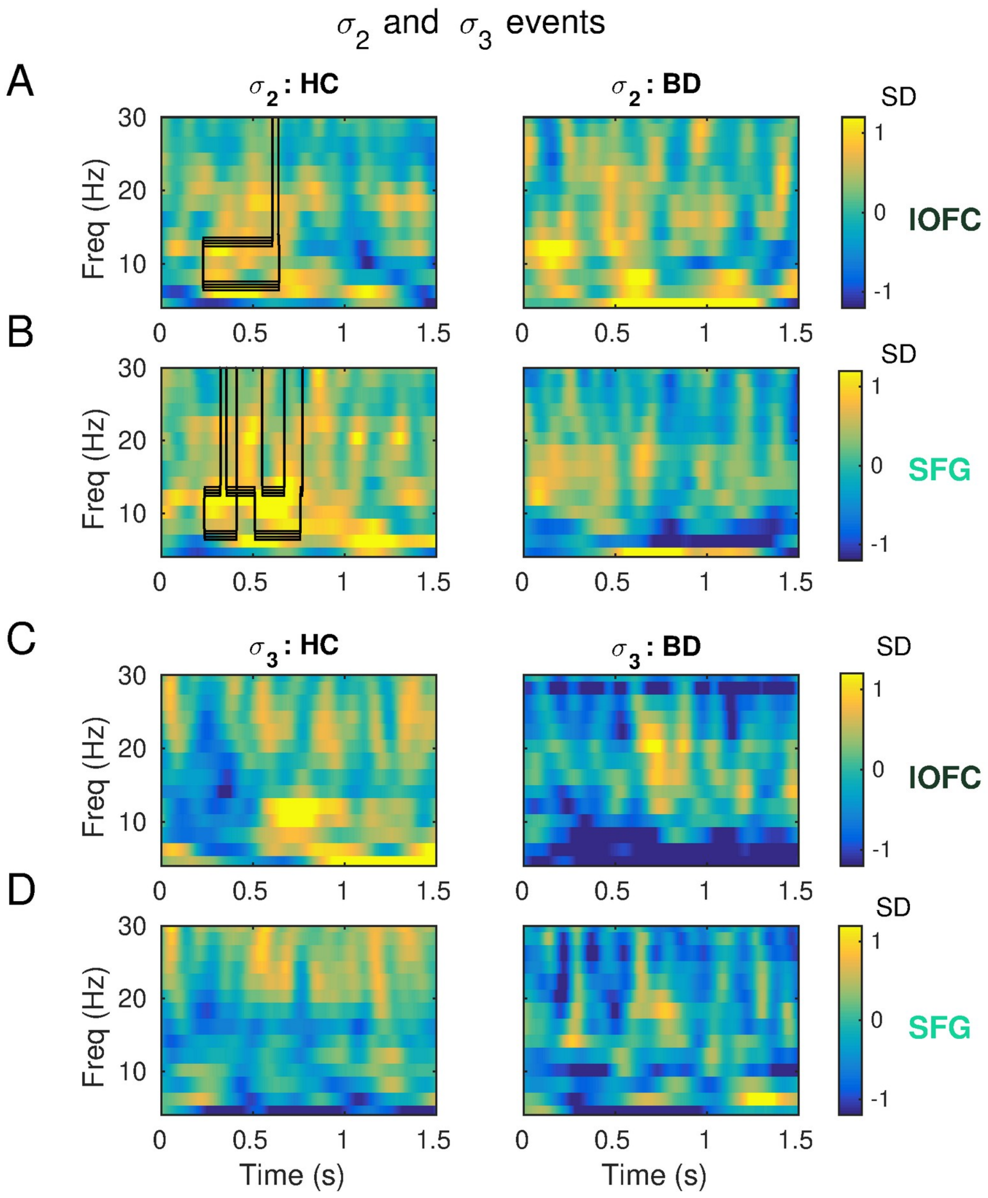

**Figure S9. Modulation of alpha and beta activity by uncertainty regressors in each group. a-d**. Same as **Figure 4**, but showing alpha and beta activity modulation by uncertainty regressors $\sigma_2$ and $\sigma_3$ in healthy control (HC, left) and bipolar disorder (BD, right) participants. For HC participants, significant increases in 8-30Hz activity were observed with estimation uncertainty, $\sigma_2$, in regions cACC, rACC, cMFG, rMFG, lOFG, M1, and SFG ($P_{FWER}$ = 0.0130, effects in regions lOFG and SFG are illustrated in panels **a-b**. No significant within-subject effects were found in BD, and there were no significant between-group differences after FWER control either (*P* = 0.1928, 0.2358 for $\sigma_2$ and $\sigma_3$, respectively). **c-d.** Regressor $\sigma_3$ was not associated with any significant within-subject or between-group differences after FWER control.

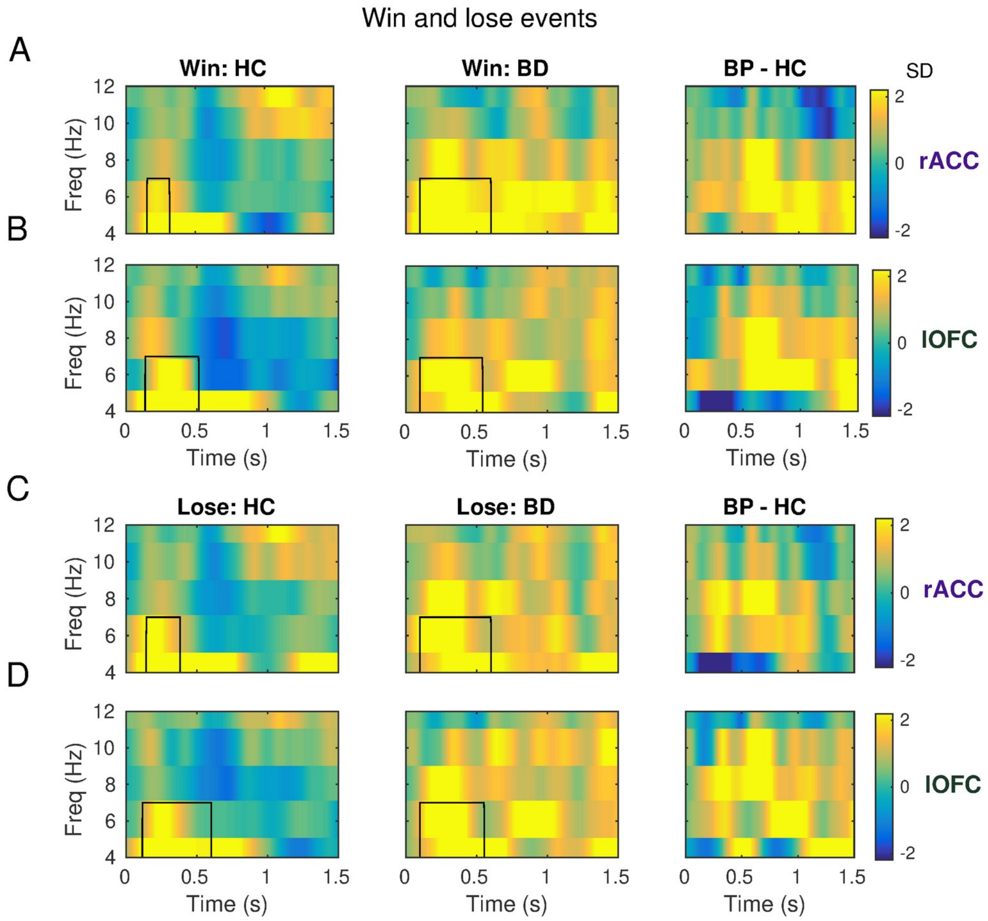

**Figure S10. Modulation of theta activity by win and lose events in bipolar and healthy control groups. a-d**. Same as **Figure 4** but for the discrete win and lose events. As expected, the discrete win (**ab**) and lose (**cd**) regressors elicited robust and significant increases in theta activity in both participant groups from 0.1 to 0.4 s ($P_{FWER}$ = 0.0240, 0.0010 for win events and HC and BD, respectively; $P_{FWER}$ = 0.0070, 0.0010 for lose events in each group). These effects were more pronounced in the caudal and rostral ACC, caudal and rostral MFG, lOFC, and mOFC, and extended for at least 200 ms (one cycle at 5 Hz) in these regions. The effects lasted longer in the bipolar group, extending from approximately 0.1 to 0.6 s. Bipolar and healthy control groups did not differ significantly with regard to the increase in theta activity in response to win or lose events (*P* = 0.1808, 0.3397 for win and lose regressors, respectively).

### Changes in Raw Power Spectral Density During the Inter-Trial Interval

The convolution modelling results from outcome processing revealed that BD patients exhibited a less pronounced suppression in the alpha and beta bands when encoding pwPEs. In a **post-hoc exploratory analysis**, we aimed to determine whether this attenuated suppression of alpha and beta activity might be associated with a smaller dynamic range of spectral power in these frequency bands in BD during task performance, namely, a reduced alpha and beta raw power.

To investigate this, we analysed the raw power spectral density (PSD, in fT) during the inter-trial intervals (ITI), a period when participants were at rest, awaiting the next stimulus to make a new choice and perform the corresponding action sequence. We extracted ITI epochs between 1.7 and 0 seconds preceding the stimulus presentation and conducted LCMV as previously described. The data covariance matrix was estimated from -1.5 to 0 s, and the noise covariance matrix from -1.7 to -1.5 s.

The analysis revealed a significant group effect on the ITI PSD, attributed to a pronounced reduction in the low beta PSD in BD compared to HC (relative suppression in the 13–20 Hz range, $P_{FDR}$ = 0.006). This effect was observed across the lateral and medial OFC, and cACC and rMFG.

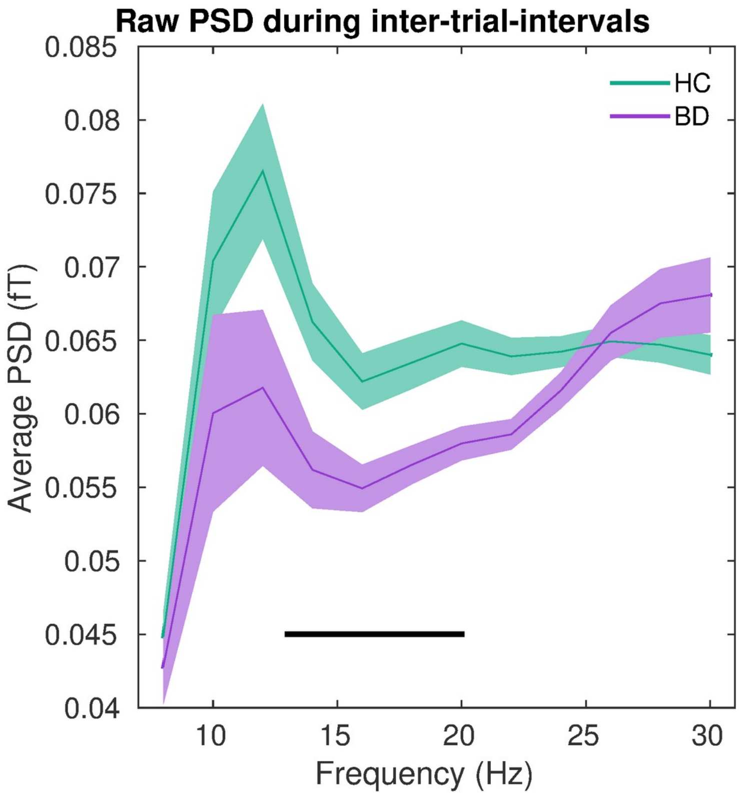

**Figure S11. Power spectral density (PSD) during inter-trial-intervals.** Grand-average of the raw PSD (in fT) during inter-trial-intervals in healthy control (HC, green) and bipolar disorder (BD, purple) participants. A significant between-group difference was obtained within 13–20 Hz ($P_{FDR}$ = 0.006). This effect emerged in most ROIs where the pwPE effect was expressed (illustrated in **Figure4** and **Figure S8**). The cluster-average in each group is represented by the continuous lines, while shaded areas denote SEM.

### Frequency-domain connectivity patterns during pwPE processing

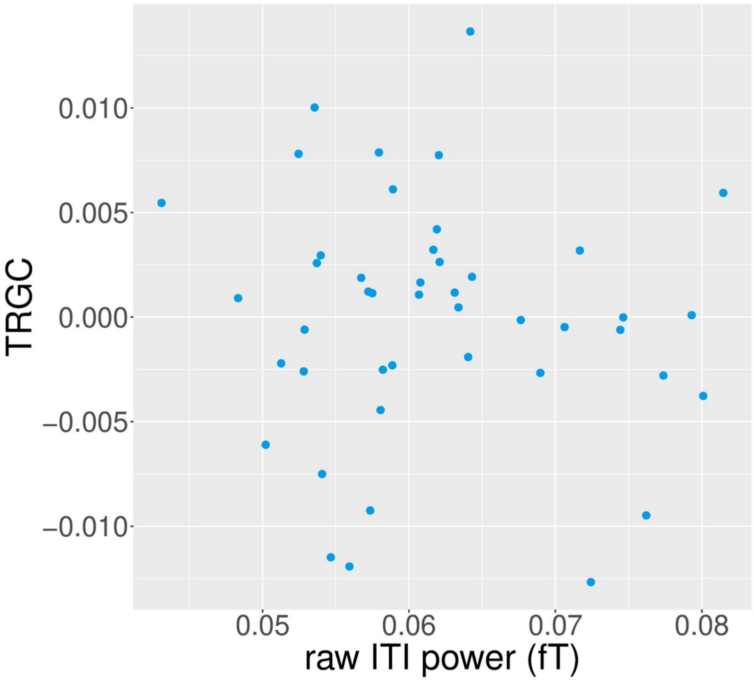

**Figure S12. Non-parametric rank correlation between average power spectral density (PSD) during inter-trial-intervals and functional connectivity metrics.** Our analyses in **Figure S11** had shown a reduction in the raw power spectral density (PSD, in fT) during inter-trial-intervals (ITI) within 13-20 Hz in BD compared to HC, denoting reduced signal-to-noise ratio (SNR) in the beta band in that time window in the bipolar group. Because SNR can influence the estimation of time-reversed Granger causality (TRGC) metrics [74], a measure of directional functional connectivity, we examined the association between beta-band ITI PSD and beta-band TRGC metrics. A BF-based Spearman rank correlation provided evidence for a lack of association between both variables (N = 49; ρ = -0.04 [-0.30, 0.24], $P$ = 0.644; $BF_{10}$ = 0.355, supporting $H_0$ based on anecdotal evidence). Thus, the main result of a significant beta-band TRGC increase in the bipolar group relative to HC (**Figure 5**) cannot be explained by an association between raw ITI PSD and functional connectivity metrics.
